# A non-genetic model of vascular shunts informs on the cellular mechanisms of formation and resolution of arteriovenous malformations

**DOI:** 10.1101/2023.08.21.554159

**Authors:** Marie Ouarné, Andreia Pena, Daniela Ramalho, Nadine V. Conchinha, Tiago Costa, Ana Figueiredo, Marta Pimentel Saraiva, Yulia Carvalho, Lenka Henao Misikova, S. Paul Oh, Cláudio A. Franco

**Affiliations:** Instituto de Medicina Molecular João Lobo Antunes, Faculdade de Medicina, Universidade de Lisboa, Lisbon, Portugal; Universidade Católica Portuguesa, Católica Medical School, Católica Biomedical Research Centre, Lisbon, Portugal; Barrow Aneurysm & AVM Research Center, Department of Translational Neuroscience, Barrow Neurological Institute, Phoenix, AZ, USA

## Abstract

Arteriovenous malformations (AVMs), a disorder characterized by direct shunts between arteries and veins, are associated with genetic mutations. However, the mechanisms leading to the transformation of a capillary into a shunt remain unclear and how shunts can be reverted into capillaries is poorly understood. Here, we report that oxygen-induced retinopathy (OIR) protocol leads to the consistent and stereotypical formation of AV shunts in non-genetically altered mice. OIR-induced AV shunts show all the canonical markers of AVMs. Genetic and pharmacological interventions demonstrated that changes in endothelial cell (EC) volume of venous origin (hypertrophic venous cells) are the initiating step promoting AV shunt formation, whilst EC proliferation or migration played minor roles. Inhibition of mTOR pathway prevents pathological increases in EC volume and significantly reduces the formation of AV shunts. Importantly, we demonstrate that ALK1 signaling cell-autonomously regulates EC volume, demonstrating that our discoveries link with hereditary hemorrhagic telangiectasia (HHT)-related AVMs. Finally, we demonstrate that a combination of EC volume control and EC migration is associated with the regression of AV shunts.

We demonstrate that an increase in the EC volume is the key mechanism driving the initial stages of AV shunt formation, leading to asymmetric capillary diameters. Based on our results, we propose a coherent and unifying timeline leading to the fast conversion of a capillary vessel into an AV shunt. Our data advocates for further investigation into the mechanisms regulating EC volume in health and disease as a way to identify therapeutic approaches to prevent and revert AVMs.

## Introduction

Arteriovenous malformations (AVMs) form as a consequence of maladaptive organisation of blood vessels. They are defined as abnormal high flow connections between an artery and a vein bypassing the capillary bed^1, 2^. Their characteristics lead to reduced tissue oxygenation and to high risks of haemorrhages and ruptures, which are often fatal when occurring in brain. The majority of brain AVMs are a consequence of sporadic events (∼95%), yet familial cases exist (∼5%)^2^. A large proportion of sporadic cases have been linked to somatic mutations in genes linked to RAS-MAPK pathway^3,4^. Congenital forms are particularly associated with hereditary haemorrhagic telangiectasia (HHT), a rare autosomal dominant genetic disorder^5, 6^. HHT is caused predominantly by *ACVRL1* and *ENG* mutations^7–9^, but can also be associated with mutations on *SMAD4*^10,11^ or *GDF2*^12–15^.

Recent advances in mouse and zebrafish animal models have provided novel insights into the mechanisms of AVM formation and progression. A common characteristic seems the cellular origin of these vascular malformations, which has been mapped to venous or capillary beds^16–18^. A second common feature is the requirement of blood flow as a driving force for AVM development^19–24^. Yet, despite these recent advances, the cellular and molecular mechanisms leading to AVM formation remain unclear. Several reports have highlighted that excessive EC proliferation is a core feature of BMP loss-of-function (LOF)-related AVM development, and interventions blocking EC proliferation such as VEGF, PI3K, AKT, or mTOR inhibitors can prevent the development of retinal AVMs in HHT mouse models^21,25–27^. In KRAS GOF, ECs showed increased cell size, ectopic sprouting, and migration properties^4,28^, and these behaviours were sensitive to MAPK inhibition but not to PI3K inhibition^28^. Alongside, cell shape changes have also been suggested to contribute to AVM development both in mouse and zebrafish models^29,30^. More recently, defective flow-migration coupling, which characterizes the ability of ECs to polarise and migrate against the blood flow direction, has been linked to HHT-like AVM models^17,21,30,31^, yet other reports showed no issues with flow-migration coupling^16,32^. Thus, to date, we lack a consensus model for AVM development and progression that could integrate all these observations and that would clarify why sporadic and familial cases develop similar vascular malformations despite arising from mutations impinging on very different signalling pathways. Here, we describe a reliable mouse model that forms AV shunts in a very predictable spatiotemporal location. This model does not rely on genetic alterations and it allows the investigation of the cellular mechanisms leading to the formation of AV shunts with very high spatiotemporal resolution. As an additional advantage, this model has the advantage to allow the study of the mechanisms of AV shunt regression, which is of special relevance to identifying new therapeutic approaches. Based on the unique features of this AVM model, we were able to propose a general model for the initiation and resolution of AVMs.

## Results

### Oxygen-induced retinopathy (OIR) model triggers transient non-genetic AV shunts

OIR is a protocol commonly used to model pathological angiogenesis, mimicking retinopathy of prematurity^33,34^. Briefly, neonatal mouse pups are exposed to hyperoxia promoting vascular regression and generating avascular retinal areas. Pups are then returned to normoxic conditions, leading to excessive and pathological neovascularization of the avascular regions. This response depends on hypoxia-driven expression of the main pro-angiogenic factor VEGFA^35^. Our protocol involves placing mouse pups at postnatal day 8 (P8) in a hyperoxia chamber until P11 (fig.1A), after which pups return to normoxia conditions. The day of return to normoxia is termed Day 0. Remarkably, we noted the rapid emergence of AV shunts in the retinal vascular network (fig.1A, B). AV shunts always form between the juxtaposed arteries and veins in the mouse retina (sup.fig.1A), and along the angiogenic border between the vascularized and avascular zone at the centre of the retina (sup.fig.1A, B). For each visible artery-vein pair, we determined the presence/absence of AV shunt and analysed the proportion of AV shunts per retina formed at specific time points. AV shunts appear starting 2 days (Day 2) after the return to normoxia (fig.1B). AV shunts diameter are variable and have a maximum mean width at Day 3 (sup.fig.1C). Interestingly, AV shunts start regressing at Day 5, and at Day 8/9 very few to none can be detected in retinas (fig.1B). This correlates with a decrease in AV shunt diameter from its peak at Day 3 (sup.fig.1C).

**Figure 1.**
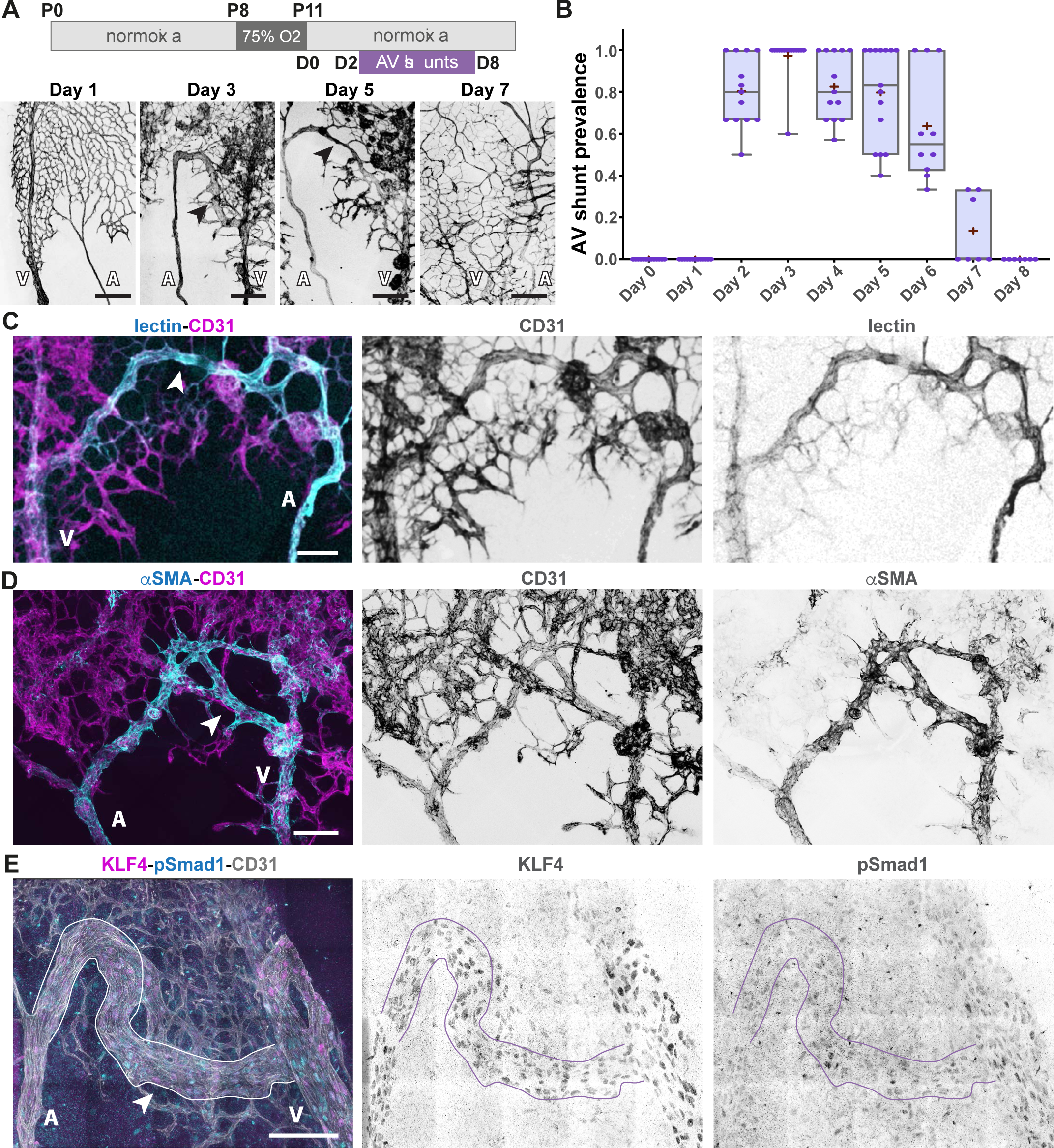
OIR protocol forms transient AV shunts independent of genetic alterations. **A,** Top panel: schematic of the experimental protocol. Bottom panel: Representative images of mouse retinas stained for CD31 (grey) on Day 1, Day 3, Day 5, and Day 7. Black arrows: AV shunt; A: artery; V: vein. Scale bar: 200 µm. **B,** Quantification of AV shunt prevalence between Day 0 and Day 8. Day 0: 51 AV sections (8 pups); Day 1: 44 AV sections (8 pups); Day 2: 56 AV sections (8 pups); Day 3: 66 AV sections (8 pups); Day 4: 55 AV sections (7 pups); Day 5: 50 AV sections (7 pups); Day 5: 50 AV sections (7 pups); Day 6: 41 AV sections (6 pups); Day 7: 30 AV sections (4 pups); Day 8: 4 AV sections (1 pup). **C,** Representative image of an AV shunt at Day 3 highlighting its perfusion status (lectin, cyan) and co-stained for ECs (CD31, magenta). Arrowhead: AV shunt; A: artery; V: vein. Scale bar: 100 µm. **D,** Representative image of smooth muscle coverage (αSMA, cyan) of an AV shunt at Day 4 co-stained for ECs (CD31, magenta). Arrowhead: AV shunt; A: artery; V: vein. Scale bar: 100 µm. **E,** Representative image of an AV shunt at Day 3 stained for ECs (CD31, grey), pSmad1 (cyan) and KLF4 (magenta). Arrowhead: AV shunt; A: artery; V: vein. Scale bar: 100 µm.

Next, we analysed OIR-induced AV shunts for characteristic features linked to genetically-driven AVMs. We observed that AV shunts are functional and carry substantial blood flow (fig.1C). This high-flow profile is further supported by high αSMA+ and desmin+ mural cell coverage (fig.1D and sup.fig.1D), and weak on NG2+ mural cells (sup.fig.1E). Additionally, high-flow AV shunts are also corroborated by high expression levels of KLF4 (fig.1E), a shear stress-responsive transcription factor^36^, which was also described as highly expressed in HHT-associated AVMs^32,37^. Together, these results demonstrate that AV shunts are high-flow vessels that become muscularized, resembling to genetically-driven AVMs. Remarkably, despite these common characteristics, the AV shunts developed in this model are not a consequence of reduced ALK1 signalling. EC nuclei in the AV shunt show normal/higher levels of Smad1 phosphorylation in comparison to capillary vessels (fig.1E), suggestive of ongoing BMP-ALK1 signalling. Overall, we identified and characterised a non-genetic model of AV shunt formation that phenocopies genetically-driven AV shunts. We propose that this model allows investigation of both the formation and the regression of AVMs with high spatiotemporal resolution.

### Non-genetic AV shunt formation is preceded by the enlargement of venules

Next, we analysed with higher temporal resolution the process of AV shunt formation. As AV shunts arise between Day 1 and Day 2, we collected retinas every 4h from timed animals between 24h and 48h. The first AV shunt start appearing at 32h. Around 36h, almost all retinas present AV shunts, with around 40% of AV segments forming a patent AV shunt (fig.2A and B). This means that in around 8h (from 28h to 36h) a large proportion of capillaries connecting AV segments converted into an AV shunt (fig.2A and B). Thus, we concluded that AV shunt formation is a progressive but rapid process. The speed of this process is similar to what is observed in Alk1-defficient mice, where AV shunts can be observed 24h post Alk1 inactivation^7–9^. To understand how such a quick conversion of capillaries into an AV shunt is possible, we studied the architecture of the vascular network during this time window (24h to 48h). As shunts preferentially develop at the limit between the avascularised and the vascularised area (sup.fig.1A and B), we characterised arteries and veins in this region. Artery and vein diameters show an increase prior to AV shunt formation. Vein and arteries when compared to diameters of vessels from animals at Day 0 immediately collected after the hyperoxia period (0h) (sup.fig.2A and B). At 24h (Day 1), the artery mean diameter is ∼121% (mean at Day 0 = 10.2 µm vs Day 1 = 12.5 µm) and the vein diameter is ∼163% (mean at Day 0 = 17.5 µm vs Day 1 = 28.5 µm) bigger than Day 0 diameters (sup.fig.2A and B). However, this change was not significant as there was a large variability between animals. This increase in vessel diameter continues over time, and by 32h changes became significant in veins and by 40h in arteries (sup.fig.2B). By 40h, when most of AV shunts have already developed, the artery mean diameter increased by ∼166% and the vein diameter by ∼224% (sup.fig.2A). Yet, remarkably, these effects were even more pronounced in second order branches of veins. The diameters of vessels connecting to arteries (arterioles) and vessels connecting to veins (venules) increase more markedly before shunt formation, when compare to parent vessels (fig.2C,D and sup.fig.2B). For instance, by 40h, the arteriole mean diameter had increased by ∼196% (mean at Day 0 = 7.0 µm vs 40h = 13.7 µm) and the venule diameter by ∼290% (mean at Day 0 = 4.5 µm vs 40h = 13.0 µm) (fig.2D and sup.fig.2B). Remarkably, the diameter of the first venule was already fully enlarged at 24h, maintaining a stable vessel width over time, whilst the other venules and arterioles increase gradually and peak at 32-40h post-normoxia (fig.2C, D and sup.fig.2B). Given that AV shunts tend to form at the first venous connection (sup.fig.1B), these results suggest that structural adaption of the first venules precedes AV shunt formation. Given that blood flow is required for AVM formation^19,20,22,23,38^, we hypothesise that OIR-dependent venule diameter increase predisposes capillary vessels to develop AV shunts by promoting unregulated blood flow between high-flow segments (proximal arteries and veins). To investigate this hypothesis, we next focused on the cellular mechanism leading to venule diameter increase at Day 1.

**Figure 2.**
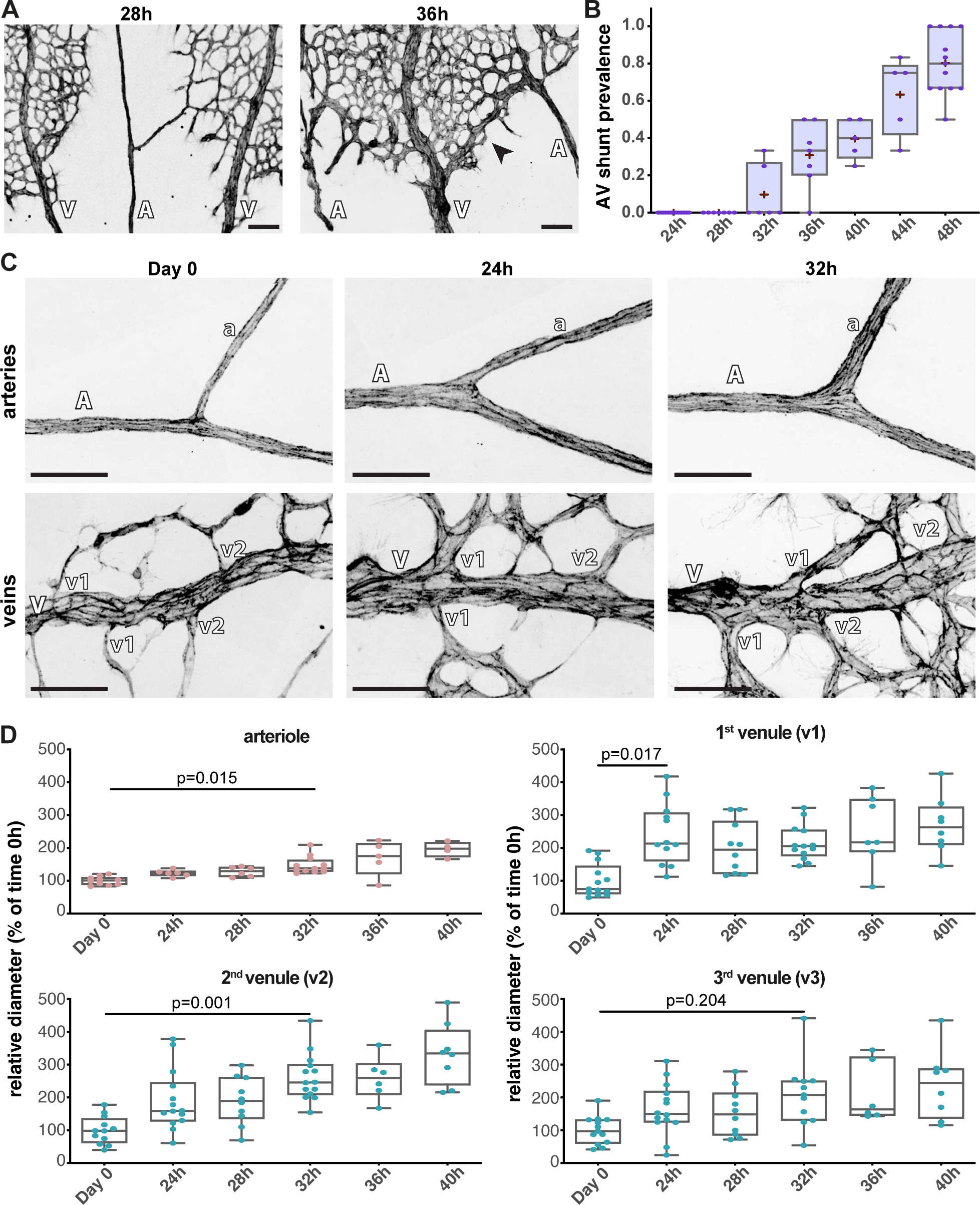
Fine time-course analysis of AV shunt formation. **A,** Representative images of mouse retinas at 28h and 36h stained for CD31 (grey). Black arrow: AV shunt; A: artery; V: vein. Scale bar: 200 µm. **B,** Quantification of total AV shunt prevalence every 4h between 24h and 48h. 24h: 44 AV sections (8 pups); 28h: 31 AV sections (7 pups); 32h: 29 AV sections (5 pups); 36h: 34 AV sections (5 pups); 40h: 23 AV sections (3 pups); 44h: 23 AV sections (3 pups); 48h: 56 AV sections (8 pups). P-values from Fisher exact t-test and Fisher post-hoc test using Benjamini & Hochberg correction for multiple comparisons. **C,** Representative images of arterial (top) and venous (bottom) second-order vessels (A: artery a: arteriole, V: vein, v: venule) at 0h, 24h, and 32h of AV shunt protocol exposed mouse retinas stained for CD31 (grey). Scale bar: 50 µm. **D,** Quantification of arteriole (top), first (bottom) and second (middle) venule normalized diameter (% of mean diameter at Day 0) between 0h and 40h. Each dot represents a second-order vessel from: Day 0 (3 pups); 24h (3 pups); 28h (2 pups); 32h (3 pups); 36h (3 pups); and 40h (2 pups). P-values from Kruskal Wallis test and Dunn post-hoc test using Benjamini & Hochberg correction for multiple comparisons.

### Endothelial proliferation nor endothelial migration are involved in OIR-induced AV shunt formation

First, we investigated EC proliferation, as it has been linked to AVM formation previously^21,25–27^, and that the OIR model is associated with neo-angiogenesis and extensive EC proliferation^34^. Indeed, we observed extensive EC proliferation in the different vessel beds prior to shunt formation, assessed by The fraction of EdU+ or pHH3+ ECs (fig.3A and B, sup.fig.3A and B). In order to assess the involvement of EC proliferation in AV shunt formation, we blocked cell proliferation using mitomycin C, a drug that inhibits DNA synthesis and cross-links DNA, effectively blocking cell cycle^39^. We treated pups with mitomycin C either at 0h or at 24h post-normoxia and collected retinas at Day 3 (fig.3C). Immunofluorescence for pHH3 or EdU demonstrates efficient abrogation of EC proliferation in mitomycin C-treated animals (fig.3C, sup.fig.3A-C). Accordingly, we observed a decrease in EC density in all vascular beds, including AV shunts, in mitomycin C-treated animals when compared to PBS-treated mice (sup.fig.3C). Yet, surprisingly, we observed no differences in the occurrence of AV shunts between PBS-treated and the mitomycin C-treated pups (fig.3D), nor a difference in shunt diameter between the two groups (fig.3E). Thus, we concluded that EC proliferation is not required for AV shunt formation or growth in our model.

**Figure 3.**
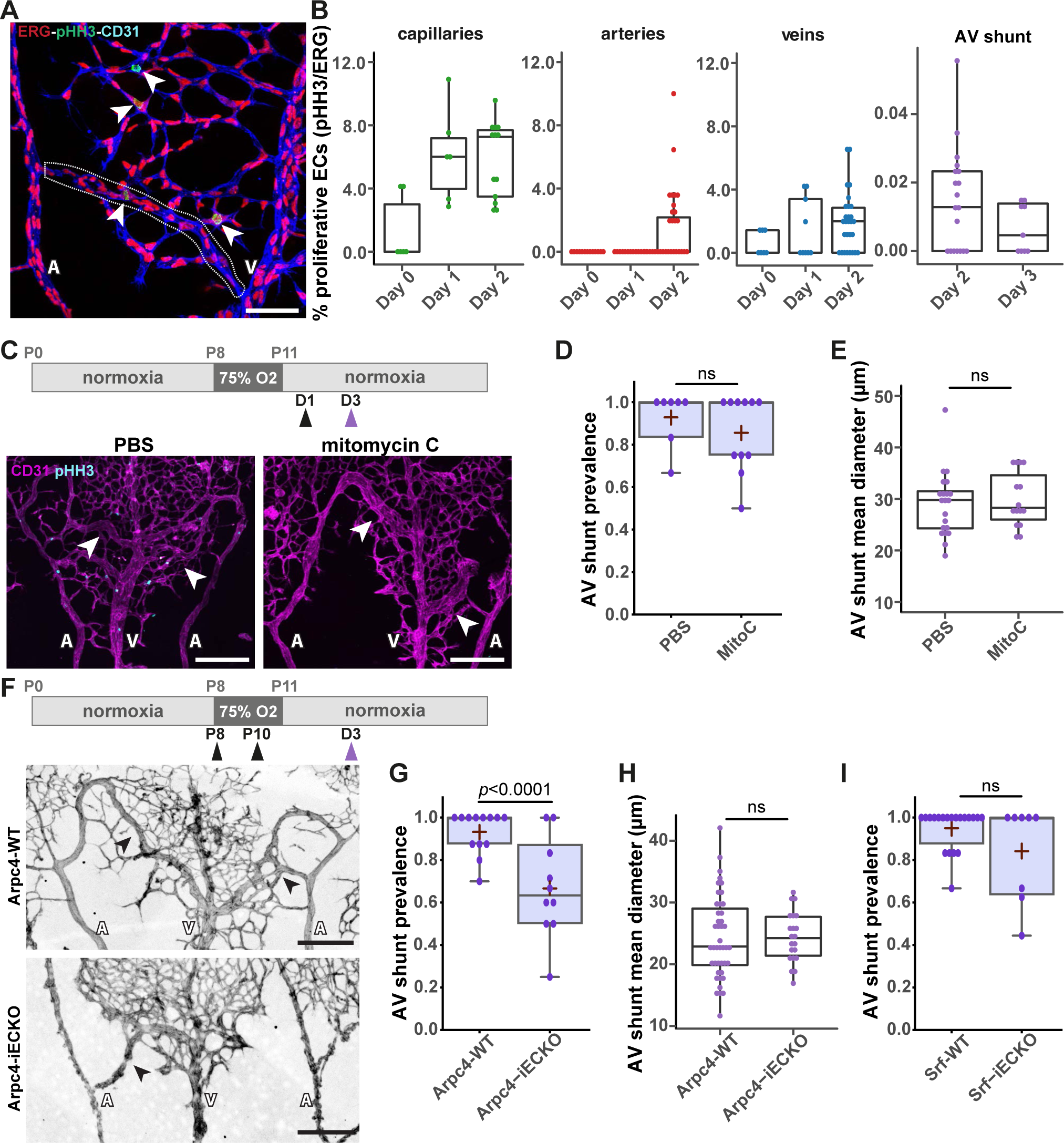
EC proliferation and migration have minor contributions to AV shunt formation. **A,** Representative image of proliferating (pHH3, green, white arrows) EC nuclei (ERG, red) within the vasculature (CD31, blue) of a Day 2 OIR mouse retina. The dotted line delineates forming AV shunt. Scale bar: 50 µm. **B,** Quantification of percentage of proliferative ECs (pHH3/ERG ratio) in capillaries, arteries and veins between Day 0 and Day 2. Each dot represents a vessel from Day 0 (4 pups); Day 1 (5 pups); and Day 2 (14 pups) retinas. P-values from Kruskal Wallis test and Dunn post-hoc test using Benjamini & Hochberg correction for multiple comparisons. **C,** Top panel: schematic of mitomycin C treatment. Black arrow: time of vehicle or mitomycin C injection; purple arrow: time of collection. Bottom panel: representative images of retinas at Day 3 treated with PBS or mitomycin C stained for ECs (CD31, magenta) and proliferative cells (pHH3, cyan). White arrows: AV shunts; A: artery; V: vein. Scale bar: 200 µm. **D,** Quantification of AV shunt prevalence at Day 3 in PBS (33 AV sections, 5 pups) and mitomycin C (46 AV sections, 7 pups) treated retinas. P-value from Mann-Whitney test. **E,** Quantification of AV shunt mean diameter at Day 3 in PBS (21 AV sections, 3 pups) and mitomycin C (14 AV sections, 2 pups) treated retinas. P-value from Mann-Whitney test. **F,** Top panel: schematic of the experimental protocol using Arpc4/Srf mouse strains. Black arrow: tamoxifen injection; purple arrow: time of collection. Representative images of Arpc4-WT and Arpc4-iECKO retinas on Day 3 stained for ECs (CD31, grey). Black arrows: AV shunts; A: artery; V: vein. Scale bar: 200 µm. **G,** Quantification of AV shunt prevalence at Day 3 in Arpc4-WT (91 AV sections, 7 pups) and Arpc4-iECKO (57 AV sections, 5 pups) retinas. P-value from Mann-Whitney test. **H,** Quantification of AV shunt mean diameter at Day 3 in Arpc4-WT (7 pups) and Arpc4-iECKO (5 pups) retinas. Each dot represents an AV shunt. P-value from Mann-Whitney test. **I,** Quantification of AV shunt prevalence at Day 3 in Srf-WT (85 AV sections, 12 pups) and Srf-iECKO (45 AV sections, 5 pups) retinas. P-value from Mann-Whitney test.

Next, we explored the role of EC migration in AV shunt development. Flow-migration coupling has been described as a player of AVM formation^17,21^. We analysed flow-migration coupling using the EC front-rear polarity GNRep mouse strain^40^ (sup.fig.4A-D). We found that at Day 0 there was a non-significant reduction in terms of polarity patterns between staged non-OIR animals (P13) and Day 0, suggesting that the OIR protocol does not affect significantly polarity patterns in arteries and veins (sup.fig.4B). Interestingly, rather than a decrease in polarity, we found a significant increase in the polarization patterns of ECs in arteries or veins, between Day 1 and Day 2, the period where AV shunts develop (sup.fig.4B). In addition, ECs in AV shunts also showed a significant polarization against the flow direction (sup.fig.4C and D). Overall, these results suggest that flow-migration coupling is not the main mechanism leading to the formation of OIR-induced AV shunts.

Next, we evaluated the contribution of EC migration in our model. We first used the endothelial-specific conditional KO of Arpc4 (*Arpc4*-iECKO) mouse line. Arpc4 is an essential subunit of the Arp2/3 complex, which creates branching actin networks that are fundamental for cell migration^41^. We previously demonstrated that Arpc4-deficient ECs showed impaired cell motility, efficiently blocking EC sprouting and EC migration in the mouse retina^41^. To avoid confounding effects, we induced *Arpc4* deletion, through tamoxifen injection, during the hyperoxia stage and collected retinas at Day 3 (fig.3F). As expected, *Arpc4* endothelial-specific deletion decreases the number of neo-angiogenic vascular sprouts during the revascularization stage (sup.fig.4E), consistent with the essential role of Arp2/3 complex in cell migration and invasion^41^. Remarkably, inhibition of cell motility led to a small, but significant, reduction in the ratio of AV shunt formation at D3 (fig.3G). Yet, inhibition of the Arp2/3 complex did not affect the diameter of existent AV shunts (fig.3H). To further confirm these results, we additionally targeted SRF in ECs. Alongside Arp2/3 complex, SRF is essential for EC migration, tip cell invasion and vessel development^42–44^. We used a similar protocol to inhibit SRF in ECs as for Arpc4, using the *Srf*-iECKO mouse line. Consistently, *Srf* endothelial-specific deletion decreased the number of neo-angiogenic vascular sprouts during the revascularization stage (sup.fig.4F). Yet, contrary to Arp2/3 complex inhibition, we did not observe a signifcant reduction in the percentage of AV shunts being formed at Day 3 in *Srf*-iECKO animals (fig.3I and sup.fig.4G). In addition, *Srf* endothelial deletion significantly decreased the diameter of AV shunts (sup.fig.4H), suggesting additional effects besides inhibition of cell migration.

Altogether, these combined results indicate that EC migration may contribute but it is not essential for AV shunt formation or development.

### EC volume changes drive AV shunt formation

Given that neither cell migration nor cell proliferation played major roles in AV shunt formation, we hypothesise that imbalances in EC distribution may cause enlargements of vessels. To examine this aspect, we decided to analyse cell density between Day 0 and Day 1 in different vascular beds, at a period preceding AV shunt formation. Within this time window, all vessel beds increase their diameters, with the first venule connection showing the highest increase in vessel diameter (fig.2D and sup.fig.2B). We quantified the number of ECs per vessel area in different vessel segments to determine local EC density (fig.4A, B and sup.fig.5A). Interestingly, the increase in vessel diameter in arteries, veins and capillaries (either on the arterial or venous side) correlated with a significant decrease in EC density (fig.4B). Remarkably, this effect was particularly strong in capillaries connecting to veins (fig.4B and sup.fig.5A). These observations suggest that the increase in vessel diameters is likely a consequence of cell volume changes rather than an increase in the number or redistribution of ECs, through proliferation or migration. This hypothesis fits with the low impact of inhibition of proliferation or cell migration on AV shunt formation (fig.3). To assess if the cell volume increases prior to shunt formation, we stochastically activated Cre recombinase, using a low dose of tamoxifen, to promote the expression of membrane-bound GFP in retinal ECs, and analysed cell shape and cell volume at the single-cell level. For this, we crossed the R26-mTmG mouse line with the Cdh5-CreERT^2^ line^45,46^, and we used previously established protocols^42,47,48^. We segmented the membrane GFP signal to generate a solid object and we analysed cell morphology and measured the total volume for each object (sup.video1). Regarding cell shapes, no significant changes in the morphology of cells could be observed, with Day 1 cells displaying similar shapes to Day 0 cells (fig.4C). Remarkably, we observed a significant increase in cell volume for ECs in all vascular beds at Day 1 (fig.4C, D and sup.fig.6A). Remarkably, the increase in cell volume is much more prominent on the venous side (∼200% increase; Day 0 mean = 1123 µm^3^ to Day 1 mean = 2332 µm^3^ for veins and Day 0 mean = 1210 µm^3^ to Day 1 mean = 2332 µm^3^ for venous capillaries) than on the arterial side (∼125% increase; Day 0 mean = 946 µm^3^ to Day 1 mean = 1227 µm^3^ for arteries and Day 0 mean = 1062 µm^3^ to Day 1 mean = 1390 µm^3^ for arterial capillaries) of the vascular tree (fig.4D and sup.fig.6A). This effect was not due to abnormal EC volume at Day 0, as cell volumes are equivalent between cells from Day 0 and non-OIR retinas in any of the vascular beds (sup.fig.6B). This effect correlates well with the bigger increase in vessel diameter on the venous side (fig.2D and sup.fig.2A). Collectively, these results point in favour that the initiation of AV shunt formation stems from an increase in EC volume on the venous side.

**Figure 4.**
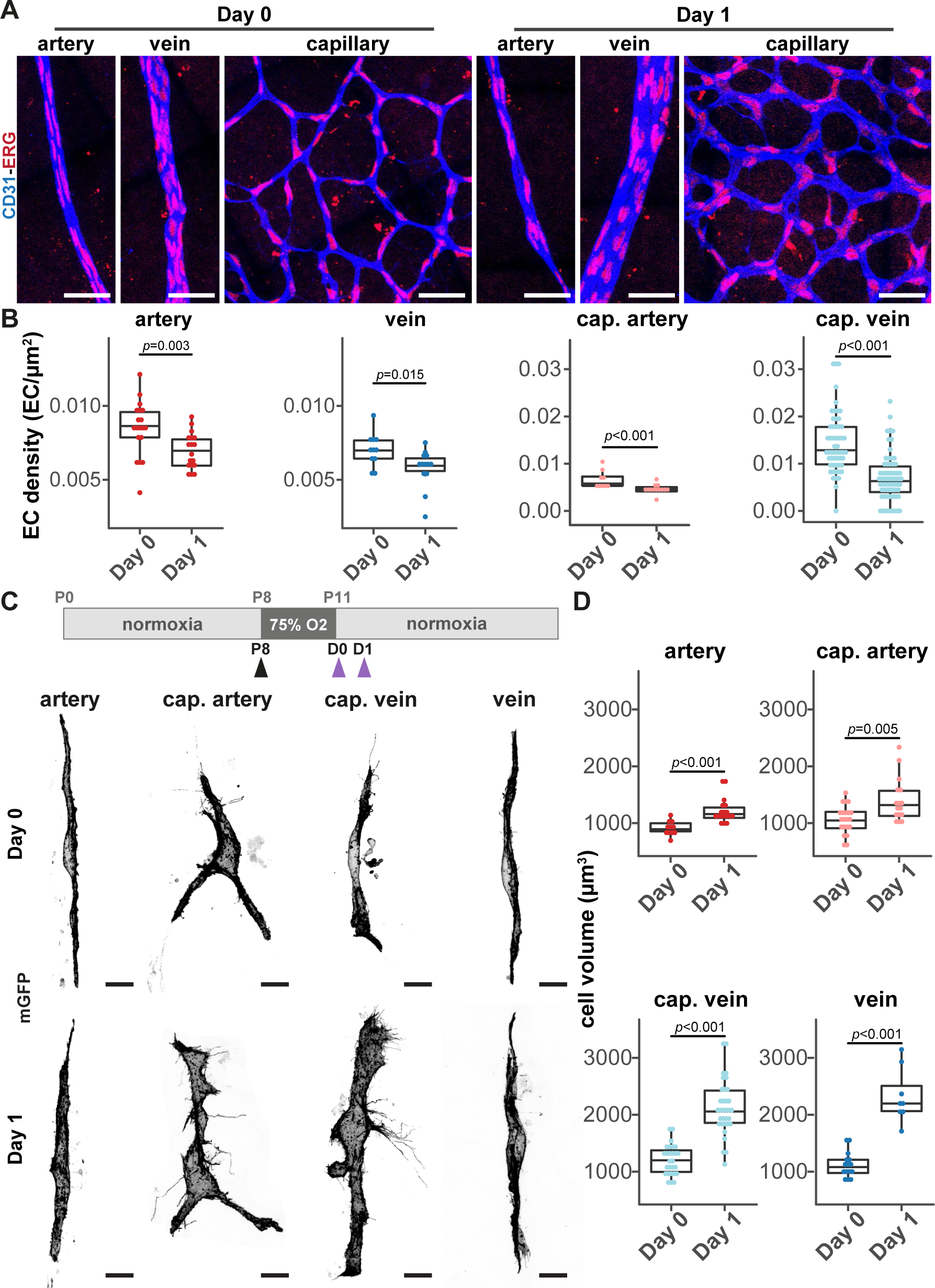
EC volume and venule diameter increases precede AV shunt development. **A,** Representative images of EC nuclei (ERG, red) distribution within artery, vein and capillary at Day 0 and Day 1 (CD31, blue). Scale bar: 50 µm. **B,** Quantification of EC density in arteries, veins, arterial capillaries and venous capillaries at Day 0 and Day 1. Each dot represents a vessel on Day 0 (4 pups) and Day 1 (5 pups). P-value from Mann-Whitney test. **C,** Top panel: schematic of experimental protocol for mosaic expression of mGFP in ECs. Black arrow: tamoxifen injection; purple arrows: time of collection. Bottom panel: representative images of single ECs (mGFP, grey) in the artery, arterial capillary, venous capillary, and vein from mouse retinas at Day 0 and Day 1. Scale bar: 10 µm. **D,** Quantification of EC volume in single cells in arteries, arterial capillaries, venous capillaries, and veins from mouse retinas at Day 0 and Day 1. Each dot represents an EC from Day 0 (3 pups) and Day 1 (3 pups). P-value from Mann-Whitney test.

**Figure 5.**
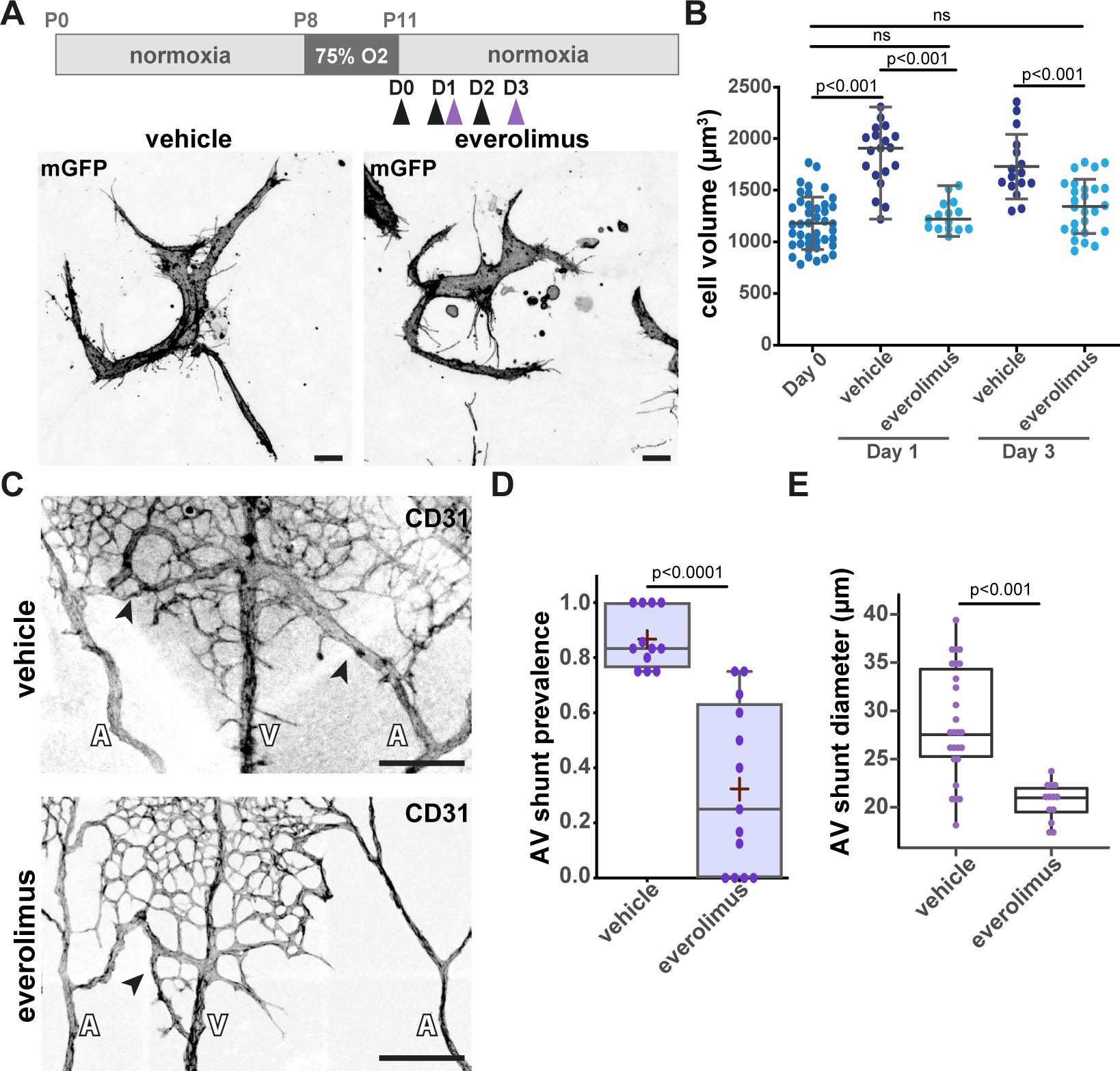
mTOR inhibition prevents EC volume increases and AV shunt formation. **A,** Top panel: schematic of AV shunt study protocol with everolimus treatment. Black arrows: times of vehicle or everolimus injections; purple arrow: time of collection. Bottom panel: representative images of single ECs (mGFP, grey) of retinas at Day 3 treated with vehicle or everolimus. Scale bar: 10 µm. **B,** Quantification of EC volume in venous cells (veins and venous capillaries) at Day 0, Day 1 and Day 3 mouse retinas treated with vehicle or everolimus. Each dot represents one EC from Day 0 (3 pups), Day 1 (3 pups) and Day 3 (3 pups). P-value from Krustal-Wallis test with Dunn’s correction for multiple comparisons. **C,** Representative images of retinas at Day 3 treated with vehicle or everolimus stained for ECs (CD31, grey). Black arrows: AV shunts; A: artery; V: vein. Scale bar: 200 µm. **D,** Quantification of AV shunt prevalence at Day 3 in vehicle (43 AV sections, 4 pups) and everolimus (54 AV sections, 5 pups) treated retinas. P-value from Mann-Whitney test**. E,** Quantification of AV shunt mean diameter at Day 3 in vehicle (4 pups) and everolimus (5 pups) treated retinas. Each dot represents an AV shunt. P-value from Mann-Whitney test.

To confirm this hypothesis, we used pharmacological inhibitors to block key metabolic pathways known to be involved in angiogenesis and control of cell volume. First, we targeted glucose, as angiogenic ECs use glycolysis as the main source of energy^49^. To do so, we use 2-deoxy-D-glucose (2-DG), which acts as a competitive substrate for hexokinase, inhibiting ATP production from glucose^50^. Remarkably, no significant changes in AV shunt formation or AV shunt diameter were observed (sup.fig.6C-E). Next, we used an inhibitor of PFKFB3 (3PO), a key glycolytic enzyme, which was identified as being critical to vessel formation and a regulator of tip and stalk cell behaviour^51^. Similar to 2-DG treatment, we observed no significant changes in AV shunt formation or AV shunt diameter were observed (sup.fig.6C-E). Thus, we concluded that glycolysis does not regulate AV shunt formation. Next, we tested the inhibition of the mammalian target of rapamycin (mTOR), a key protein complex regulating cell metabolism, cell growth and cell proliferation^52^. We used everolimus, which preferentially targets mTOR complex 1 (mTORC1). Also, mTOR inhibitors were previously shown to prevent AVM in a mouse model of HHT^27^. A single dose of everolimus at Day 0 significantly decreased EC volume at Day 1, normalizing it to volumes similar to Day 0 cells. Daily injections of everolimus between Day 0 and Day 2 maintained normalization of EC volumes at Day 3 (fig.5A and B). Importantly, everolimus-induced normalization of EC cell volumes correlated with a significantly decreased in the rate of AV shunt development at Day 3 (fig.5C and D), alongside a significant decrease in shunt diameter (fig.5E), suggesting that a change in EC volume is an essential step initiating AV shunt formation in our mouse model.

Taken together, our results collectively suggest a model describing the formation of an AV shunt (fig.6A). We propose that AV shunts originate from the abnormal and asymmetric enlargement of venous vessels due to an increase in EC volume as the main cellular mechanism. This asymmetric vessel enlargement promotes unregulated flow rates in the proximal capillary bed connected to high-flow arteries and veins, which correlates with the interface with the avascular zone in the OIR model. This initial uncontrolled flow pattern self-amplifies by the conversion of a capillary vessel path into a proper AV shunt. These later events will likely involve EC migration and EC proliferation, in addition to EC volume changes.

**Figure 6.**
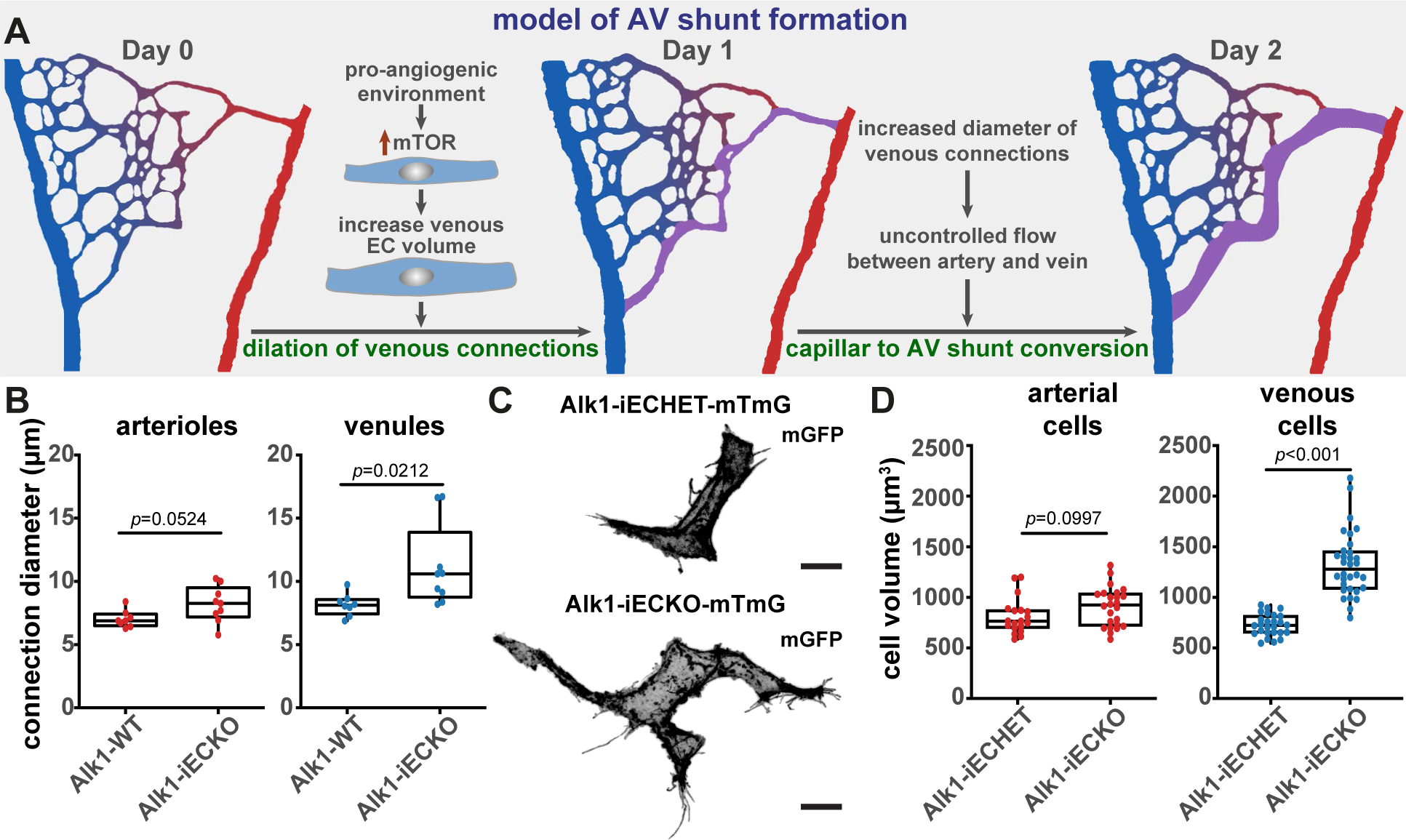
Alk1 signaling controls EC volume cell-autonomously. **A,** Model describing AV shunt formation in OIR protocol. **B,** Quantification of arteriole and venule diameter in Alk1-WT and Alk1-iECKO retinas 24h post tamoxifen injection. Each dot represents a second-order vessel. P-value from Mann-Whitney test. **C,** Representative images of ECs from a venous capillary EC (GFP, grey) of Alk1.iECHET-mTmG and Alk1-iECKO-mTmG 24h post-tamoxifen injection. Scale bar: 10 µm. **D,** Quantification of EC volume in arterial (arteries and arterial capillaries) and venous (vein and venous capillaries) vessels from Alk1.iECHET-mTmG and Alk1-iECKO-mTmG 24h post-tamoxifen injection. Each dot represents one EC from Alk1.iECHET-mTmG (2 pups) and Alk1-iECKO-mTmG (2 pups). P-value from Mann-Whitney test.

To test if the proposed model applies to genetic models of AVMs, we focused on *Alk1* LOF in ECs. Endothelial-specific deletion of *Alk1* leads to rapid (30h-36h) development of AV shunts in the mouse retina^24,53^. Concordant with our data on OIR-induced AV shunts, we observed a significant increase in vessel diameter at 24h post-recombination, a stage prior to AV shunt formation (fig.6B). This increase was more robust on the venous side, when compared to the arterial side, further corroborating the data obtained on the OIR model (fig.6B). Next, to confirm if vessel diameter increase was associated with changes in cell volume, we intercrossed the Alk1 mouse model with the R26-mTmG mouse line. Low-dose tamoxifen injection allows recombination of a few ECs (sup.fig.6F), enabling the measurement of Alk1 LOF in cell volume in the context of a WT retina (fig.6C), without generating AV shunts. Strikingly, stochastic recombination of Alk1 leads to a significant increase in EC volume at 24h post-recombination specifically in the venous regions, whilst the arterial ECs showed no significant changes (fig.6C and D). Thus, we concluded that Alk1 signalling controls EC volume in a cell-autonomous manner and that the initiation steps driving AV shunt formation share similarities between HHT-induced and OIR-induced models. This suggests that EC volume control, and concomitant flow pattern deregulation, is a key mechanistic step leading to AVM development in HHT.

### AV shunt regression is dependent on endothelial flow-migration coupling and cell volume changes but not on EC apoptosis

AV shunt regression is of particular clinical relevance yet very little is known about the underlying cellular and molecular mechanisms. Thus, we took advantage that OIR-induced AV shunts are not stable to investigate how AV shunts resolve. AV shunts start regressing at D5, and by D8 no AV shunts can be detected (fig.1B). We hypothesise that shunt resolution could be the outcome of one or a combination of several mechanisms, including EC apoptosis, cell migration or cell volume changes. First, we investigated apoptosis, which could lead to a reduction in the number of ECs in a vessel segment leading to a decrease in its diameter. Active-caspase 3 staining during the resolution phase highlights the existence of very few apoptotic ECs during this stage (sup.fig.7A), and thus we excluded apoptosis as a mechanism for AV shunt resolution.

Next, we evaluated if cell dispersion through cell migration could explain AV shunt resolution. During this stage, we observed that shunt regression coincides with the neo-vascularization of the avascular area (fig.7A and B). This leads to an increase in the number of connections between the AV shunt and the parent arteries and veins and an increase in the vascular density of the neo-capillary network (fig.7A, B and sup.fig.7B). Given that flow-migration coupling is essential for vascular remodelling and network optimisation^54–57^, we hypothesise that redistribution of blood flow through the new vascular segments could reroute EC migration paths. This may decrease the number of ECs moving into AV shunts, and therefore contribute to their normalisation. As the majority of neo-vessels connect AV shunts with the adjacent veins (fig.7A and B), we predicted that this region may show the first signs of AV shunt normalisation. To validate this hypothesis, we first assessed vascular perfusion of neo-vascular networks using intracardiac lectin injections. We observed a significant increase in the number of perfused vessel branches from both the vein and the AV shunt, from Day 2 to Day 6 (fig.7C and D), suggesting progressive blood flow redistribution from AV shunts towards newly formed vascular beds. This coincided with an increase in the overall perfusion of the neovascular area (sup.fig.7C). AV shunts start thinning and become less discernible at the venous side (fig.7A and C), where the connection between the AV shunt and the draining vein becomes more entangled. Remarkably, quantification of AV shunt diameters from the arterial and venous sides showed a preferential decrease in diameter from the venous side starting at Day 4, which precedes AV shunt regression (fig.7E). Overall, these results strongly suggest that AV shunt regress due to changes in flow distribution and EC rerouting, occurring first on the venous side.

**Figure 7.**
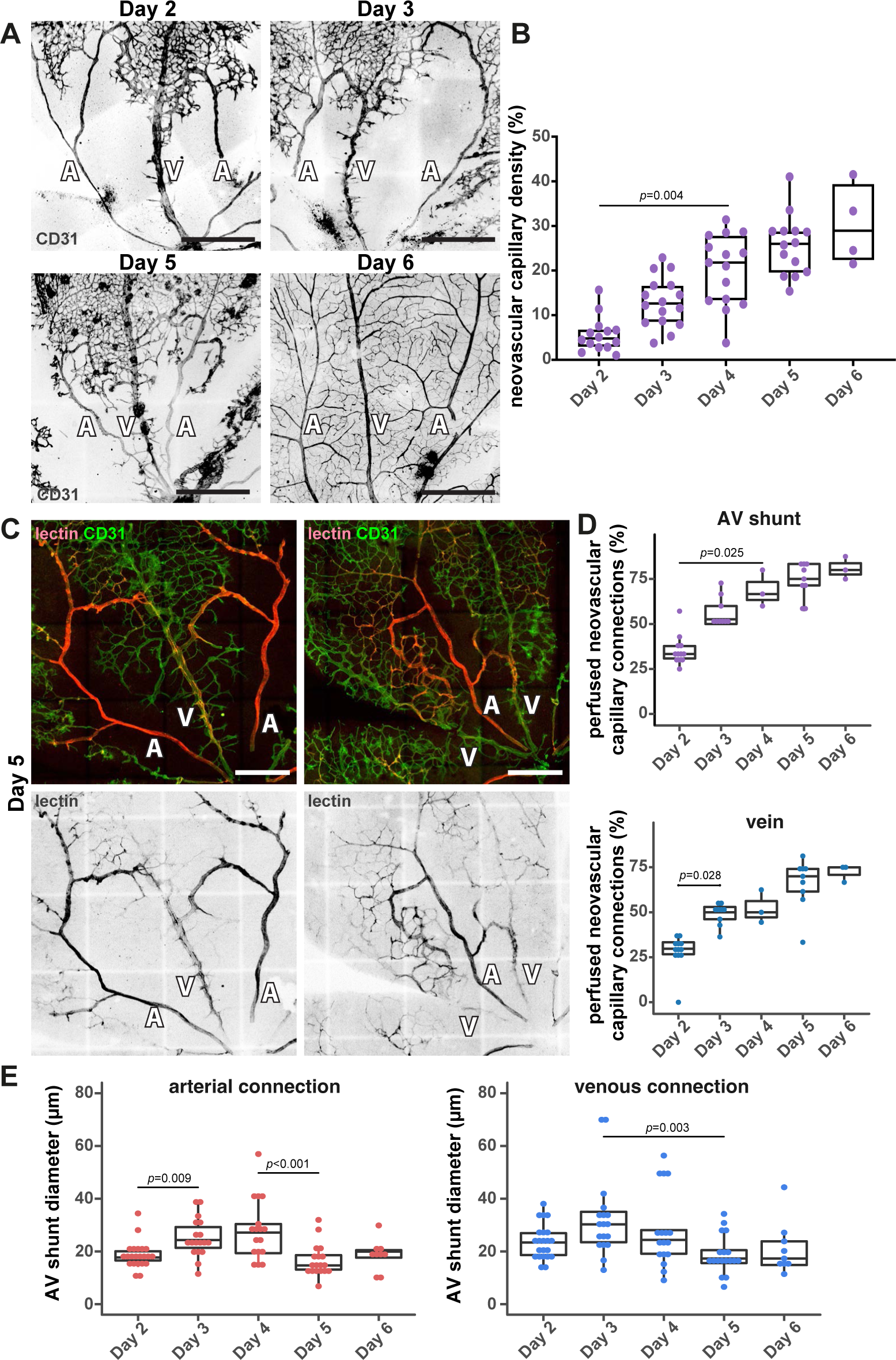
AV shunt regression correlates with perfusion of the neo-capillaries. **A,** Representative images of mouse retinas stained for CD31 (grey) at Day 2, Day 3, Day 5, and Day 6. A: artery; V: vein. Scale bar: 500 µm. **B,** Quantification of neovascular capillary density between Day 2 and Day 6. Each dot represents an AV shunt proximal region from Day 2 (4 pups); Day 3 (3 pups); Day 4 (3 pups); Day 5 (4 pups); and Day 6 (2 pups). P-values from Kruskal-Wallis test and Dunn post-hoc test using Benjamini & Hochberg correction for multiple comparisons. **C,** Representative images of Day 5 mouse retinas perfused with lectin (red) and co-stained for ECs (CD31, green). A: artery; V: vein. Scale bar: 250 µm. **D,** Quantification of perfused neovascular capillary connections to AV shunt (top) and associated vein (bottom) between Day 2 and Day 6. Each dot represents an AV shunt or a vein from Day 2 (6 retinas); Day 3 (4 retinas); Day 4 (1 retina); Day 5 (5 retinas); and Day 6 (3 retinas). P-values from Kruskal-Wallis test with Dunn’s correction for multiple comparisons. **E,** Quantification of AV shunt diameter on the first 50 µm connected to the corresponding artery (left) or vein (right) between Day 2 and Day 6 mouse retinas. Each dot represents an AV shunt from Day 2 (4 pups); Day 3 (3 pups); Day 4 (3 pups); Day 5 (4 pups); and Day 6 (2 pups). P-values from Kruskal-Wallis test and Dunn post-hoc test using Benjamini & Hochberg correction for multiple comparisons.

To tackle the importance of the blood flow redistribution and EC migration-flow coupling in AV shunt regression, we took advantage of the *Arpc4*-iECKO and *Srf*-iECKO mouse models to inhibit EC migration and invasion. To avoid any impact on the formation of AV shunts, we induced *Arpc4* or *Srf* deletion after return to normoxia. As expected, *Arpc4*-iECKO pups showed a significant reduction in the neo-angiogenic sprouts and a significant reduction in the vascularization of the avascular region (sup.fig.6D and E). Remarkably, at Day 7, *Arpc4*-iECKO pups preserved AV shunts, contrary to control pups, where AV shunts almost completely regressed (fig.8A and B). A similar trend was found when inhibiting SRF function in ECs. *Srf*-iECKO pups also showed a significant reduction in the number of neo-angiogenic sprouts (sup.fig.7F), alongside the maintenance of AV shunts at Day 9 (fig.8C and D). Altogether, we concluded that formation of a neo-vascular network is essential for AV shunt regression.

**Figure 8.**
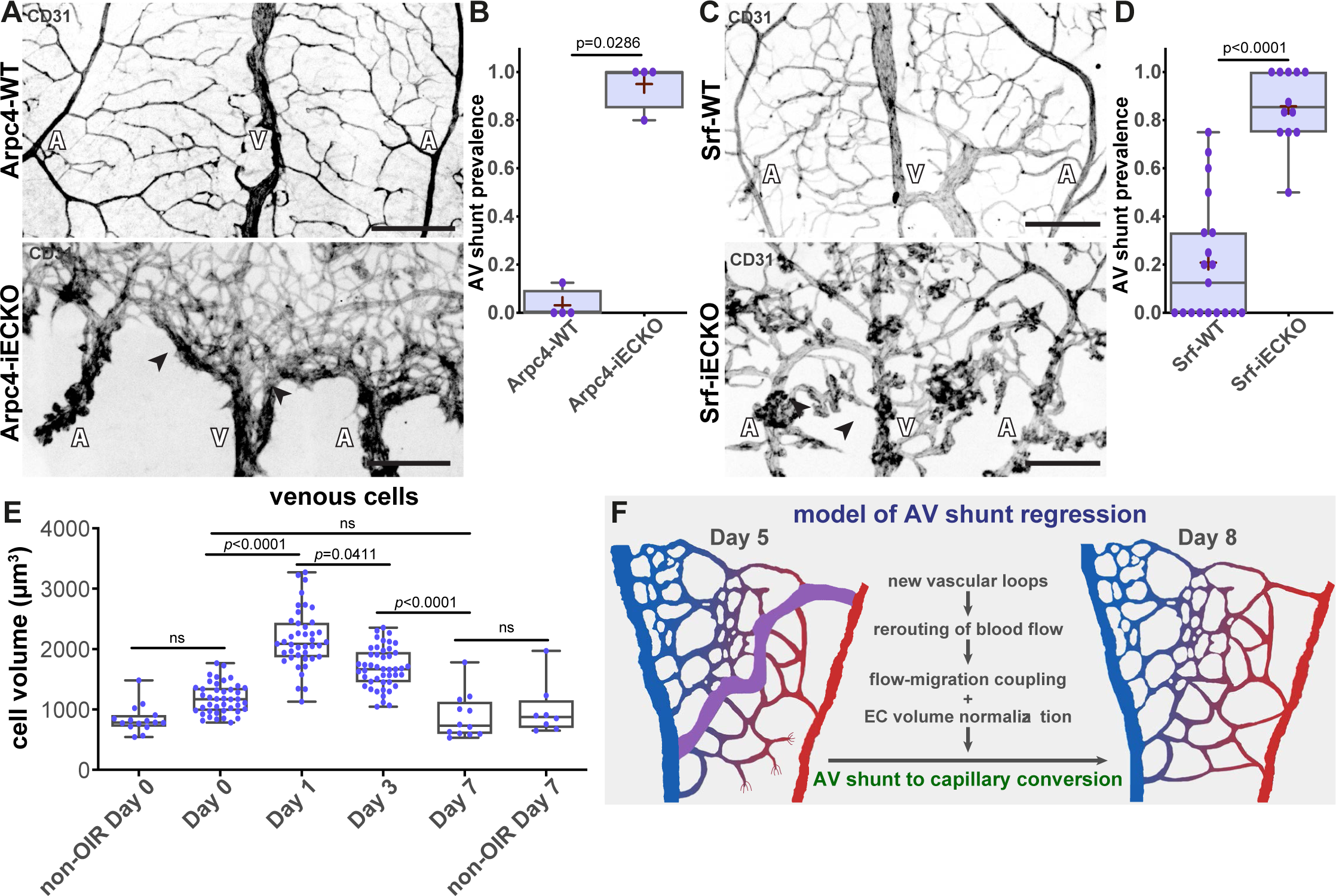
Inhibition of neo-vascular capillaries prevents AV shunt regression. **A,** Representative images of Arpc4-WT (top) and Arpc4-iECKO (bottom) retinas at Day 7 stained for vascular network (CD31, grey). Black arrows: AV shunts; A: artery; V: vein. Scale bar: 200 µm. **B,** Quantification of AV shunt prevalence at Day 7 in Arpc4-WT (29 AV sections) and Arpc4-iECKO (16 AV sections) retinas. P-value from Mann-Whitney test. **C,** Representative images of Srf-WT (top) and Srf-iECKO (bottom) retinas at Day 9 stained for vascular network (CD31, grey). Black arrows: AV shunts; A: artery; V: vein. Scale bar: 200 µm. **D,** Quantification of AV shunt prevalence at Day 9 in Srf-WT (86 AV sections) and Srf-iECKO (63 AV sections,) retinas. P-value from Mann-Whitney test. **E,** Quantification of venous (vein and venous capillaries) EC volume at Day 0, Day 1, Day 3, and Day 7 and of non-OIR mouse retinas corresponding to time points Day 0 (P11) and Day 7 (P18). Each dot represents one EC from Day 0 (3 pups,); Day 1 (3 pups); Day 3 (3 pups); Day 7 (2 pups); non-OIR Day 0 (2 pups); and non-OIR Day 7 (2 pups). P-values from Kruskal-Wallis test with Dunn’s correction for multiple comparisons. **F,** Model describing AV shunt regression in OIR protocol.

Finally, we analysed EC volume. Given that an increase in EC volume is the driving force for AV shunt formation (fig.4 and 5), we examined if the regression of AV shunts could correlate with a decrease in EC volume. To do so, we employed a similar approach as for AV shunt formation, but we induced stochastic recombination of the reporter line at Day 0 instead. The analysis of ECs in each vascular bed demonstrated normalization of cell volume in arterial and venous cells, obtaining volumes comparable to Day 0 (fig.8E). Thus, taken together, our data points towards a model where AV shunts resolve by a combination of EC dispersion through flow-migration coupling and EC volume normalization (fig.8F). Both mechanisms may cooperatively promote the reversion of abnormally formed AV shunts into capillary vessels of normal diameter.

## Discussion

In this work, we unravelled a new non-genetic mouse model to study AV shunt formation and regression with high spatiotemporal resolution. Through genetic and pharmacological interventions, we mechanistically demonstrated that EC volume control, rather than EC proliferation, is a key step in the formation of AV shunts, whilst a combination of cell volume control and EC flow-migration coupling is associated with the regression of these vascular malformations.

Based on our results, we propose a coherent and unifying timeline leading to the fast (24h) conversion of a capillary vessel into an AV shunt. A first trigger, either genetic mutation or specific environmental conditions, leads to a substantial increase in the volume of ECs and a concomitant increase in vessel calibre. Remarkably, our detailed analysis showed that the early changes driving both AV shunt development and regression are preferentially located in the venous compartment (veins and venous capillaries). This is in agreement with previous studies on HHT-associated AVMs that pointed towards ECs in capillaries or veins as the main cellular origin of vascular shunts^16–18,21^. This increase in cell volume is fuelled, at least partially, via the mTOR pathway, and it leads to the expansion of the capillary diameter. In turn, the expansion in vessel diameter decreases flow resistance leading to an increase in flow rates in the vessels prone to be converted into shunts. Increased levels of blood flow through these dilated vessels further expand the vessel lumen through a combination of EC migration, flow-migration coupling and EC proliferation. Recruitment or transdifferentiation of mural cells promotes the consolidation of the high-flow shunt. In addition, we showed that AV shunts can gradually regress through a combination of flow-migration coupling-induced remodelling and cell volume changes. The regression relies on the intrinsic capacity of ECs to migrate and rearrange within the vascular network in order to resolve maladaptive vessel configurations, an essential behaviour that we refer to as vascular plasticity^57^. In this context, the establishment of new vessel connections, which create new flow routes is a prerequisite for AV shunt regression, and blockage of vascular plasticity can sustain environmentally-driven AV shunts.

This detailed description of the initial steps in the formation and resolution of AVMs raises further questions. What is the relative contribution of EC volume control in shunt regression in relation to EC migration and EC redistribution? A large contribution of EC volume normalization would point towards a common cellular process involved in the genesis and resolution of these vascular anomalies, whilst a low contribution would indicate fundamentally different mechanisms of the two biological phenomena. The latter could indicate that known mutations driving AV shunts may impair two distinct cellular processes, one leading to EC volume increases and another disrupting vascular plasticity, which will promote the formation and, at the same time, prevent the mechanisms of regression.

One additional key question resides in the molecular mechanisms of EC volume control. So far, this question has raised very limited attention in the field of vascular biology. Yet, previous connections between cell size have been reported to be associated with AVM formation. For instance, HHT-driven mutations have been shown to lead to bigger EC sizes in zebrafish and mice^29,30^. Moreover, KRAS-activating mutations have recently been identified as the main driver of sporadic brain AVMs and they were also associated with increases in EC volume^4,28^. However, constitutively active Notch4 also gives rise to AVMs with increased capillary diameters, yet cell volume changes have not been reported so far^22,38,58^. How these pathways regulate cell volume remains to be elucidated. Generally, short timescale cell volume control is achieved through osmolarity control, mainly via ion channels^59,60^. Longer timescale cell volume control has been mostly studied in the context of cell cycle and has been associated with several pathways promoting anabolism, such as the mTOR pathway^52,61^, MYC signalling^62,63^, the YAP/TAZ pathway^64–66^, and more recently cell mechanics^67,68^. Even if we cannot exclude the impact of osmolarity effects and short timescale fluctuations on cell volume, the significant normalisation of EC volumes upon mTOR pathway inhibition with everolimus treatment (fig.5), and concomitant impact on AV shunt formation, strongly suggests that anabolic activity is a key fundamental step in pathological EC volume control. How AVM-associated pathways regulate anabolism may differ according to the associated mutations. For instance, KRAS activating mutations rely on MEK activity rather than on AKT/PI3K signalling, an upstream regulator of mTOR activity^4,28^, whilst ALK1, ENG or SMAD4 LOF mutations showed sensitivity to AKT/PI3K signalling inhibitors^21,25,26^. In this regard, OIR-induced AV shunts are more closely related to HHT-associated lesions rather than to KRAS-induced AVMs. Remarkably, BMP pathway LOF mutations require pro-angiogenic environments to induce AVM formation whilst KRAS activating mutations are able to promote AVM development in quiescent endothelium^24,26,28,69^, which further points towards a closer mechanistic relationship between OIR-induced AV shunts with HHT-associated AVMs.

Despite the strong evidence of cell volume as a key mechanism driving AVM formation, how EC volume-dependent lumen enlargement feedbacks into flow dysregulation that promotes capillary-to-shunt conversion remains largely obscure. Through a rheological perspective, differential resistance of capillary vessel segments would explain preferential shunting of flow through enlarged vessels, yet vascular cells have evolved numerous mechanisms tightly controlling blood flow, with a particular emphasis on mural cells^70^. Thus, it is likely that additional mechanisms related to mural cell activity may be affected in our model which further promotes AV shunt development. Interestingly, recent reports have also linked mural cell function and AVM formation in animal models^71–73^. Yet, to our perspective, dysfunction of mural cell-dependent flow control is rather a facilitator rather than a driver, and pre-requires an imbalance in EC volume as an initiating step.

Finally, our work also establishes a solid model to investigate AVM regression. How and why genetically driven AVMs do not regress is a key open question. Recently, thalidomide treatment has shown promising effects on the regression of AVMs in patients with a severely symptomatic AVM that is refractory to conventional therapies^74,75^. Yet, the molecular mechanisms of the action of this broad-spectrum drug remain unclear. Taking our results into consideration, we can propose that thalidomide may either promote cell volume normalization and/or efficient flow-migration coupling-induced remodelling.

Moreover, our novel insights into AV shunt regression also open the perspective of a novel class of mutations that might be associated with human AVMs. As AV shunts can naturally occur in genetically-competent individuals, mutations impacting AV shunt resolution mechanisms, rather than AV shunt formation mechanisms, could promote the stabilisation and growth of those lesions by a lack of capacity to resolve them. Given that mutations in Srf and Arp2/3 complex limit new sprout formation and AV shunt regression, we predict that mutations impacting EC motility and sprouting when associated with naturally occurring shunts may lead to AVMs.

In conclusion, we demonstrated that EC volume is the key mechanism driving AVM formation, and it seems transversal to genetic and non-genetic AVM mouse models. Our data strongly underline the necessity to further investigate the mechanisms regulating EC volume in health and disease as a way to identify therapeutic approaches to prevent and revert AVMs.

## Materials and methods

### Mice

All animal experiments carried out in this work were performed in compliance with the relevant laws and guidelines that apply to the Instituto de Medicina Molecular (iMM) – João Lobo Antunes, Faculty of Medicine, University of Lisbon, Portugal. Animal procedures were performed under the Direção-Geral da Alimentação e Veterinária (DGAV) project licenses 012092/2016 and 017722/2021.

Mice were maintained at the Instituto de Medicina Molecular (iMM) under standard husbandry conditions (under specific pathogen-free conditions and kept in individually ventilated cages) and under national regulations.

The following transgenic mouse strains were used in this study: GNrep^40^, *Arpc4* floxed^76^, *Srf* floxed^77^, *Alk1* floxed^24^, and R26-mTmG^45^ mice. The different strains were crossed with Cdh5(PAC)-CreERT2 strain^46^ or Pdgfb-CreERT^78^ to obtain the desired genotypes. Cre-negative littermates were used as controls in KO strain experiments. Both males and females were used, without distinction. Animals were sacrificed at different endpoints and the eyeballs were collected.

### Treatments

4-hydroxytamoxifen (H6278, Sigma-Aldrich) was injected intraperitoneally (IP) (20 μg/g) at different ages depending on the mouse strain and studied AV shunt stage. KO strains were injected at post-natal day 8 (P8) and P10 to study AV shunt formation or at Day 1 (P12) and Day 2 (P13) to study AV shunt resolution. GNrep mice were injected at P1 and P3 to trigger reporter expression. To trigger mosaic recombination in R26mTmG, a low dose of 4-hydroxytamoxifen (0.4 μg/g) was injected IP only once at P8 or P14. Non-OIR pups were injected with the same low dose of 4-hydroxytamoxifen tamoxifen three days before the day of collection.

To block cell proliferation during shunt formation, mitomycin C (SC-3514B, ChemCruz) was injected IP (10 μg/g) at Day 0 or Day 1 and pups were collected at Day 3. PBS was injected in control pups.

To quantify EdU+ cells *in vivo*, a stock of 50mg 5-ethynyl-2-deoxyuridine (EdU) (Alfagene, A10044) was diluted in 5mL of PBS to make a working solution (10mg/mL). EdU solution was injected intraperitoneally (200mg/kg) 4 hours before the animals were sacrificed. Retinas were isolated and fixed as previously described, and the EdU-positive cells were detected according to the user manual of the Click-iT EdU Alexa Fluor 555 Imaging Kit (Invitrogen, C10338).

To affect glucose metabolism during shunt formation, PFKFB3 inhibitor (3PO; 50 μg/g) (525330, Merck Life Sciences) or 2-deoxy-glucose (2-DG; 500 μg/g) (25972, Merck Life Sciences) were injected IP at Day 0, Day 1, and Day 2 and pups were collected at D3. For both treatments, PBS was injected in control pups. mTOR pathway was inhibited using everolimus (13,5 μg/g) (73124, Stemcell) in peanut oil at Day 0, Day 1 and Day 2 to study shunt formation. Ethanol (vehicle) diluted in corn oil was injected to control pups.

Vascular perfusion was assessed by injecting 10 μl of DyLight-649 conjugated Lycopersicon Esculentum (Tomato) lectin (DL-1178-1, Vector Laboratories) intracardially (1mg/mL) in anesthetise pups at let circulate for a minimum of 5 min time before mouse sacrifice and eye collection.

### Hyperoxia chamber protocol

P8 pups and their nursing mothers were housed in a Biospherix A-Chamber (Biospherix) equipped with a ProOx 110 oxygen controller (Biospherix). In the chamber, the animals were exposed to 75% oxygen level from P8 until they return to normal room air conditions at P11, also termed as Day 0. Pups were sacrificed at the time points. Eyes were collected and fixed with 2% PFA (15710, Electron Microscopy Sciences) in PBS for 4 hours at 4°C.

### Immunofluorescence on mouse retinas

Retinas were dissected in PBS and stained according to previously established protocols^79,80^. Briefly, retinas were incubated on a rocking platform for 2 hours at room temperature (RT) in Claudio’s blocking buffer (CBB) consisting of 1% FBS (LTID 10500-064, Thermo Fisher), 3% BSA (MB04602, Nzytech), 0.5% Triton X100 (T8787, Sigma Aldrich), 0.01% sodium deoxycholate (30970, Sigma Aldrich), 0.02% sodium azide (S2002, Sigma Aldrich) in PBS, pH = 7.4. Primary antibodies (see Table 1) were incubated in 1:1 CBB/PBS overnight on a rocking platform at 4°C. Afterwards, retinas were washed 3 times 30 minutes with PBS 0.1% triton X-100 (X100, Sigma Aldrich). Secondary antibodies (see Table 1) were incubated in 1:1 CBB/PBS overnight on a rocking platform overnight in the dark at 4°C. Then retinas were washed 3 times 30 minutes with PBS 0.1% Tween and mounted on slides using Vectashield mounting medium (H-1000, VectorLabs).

**Table 1:** Antibodies.

| Antibody | Conjugation | Manufacturer | Reference | Dilution |
| --- | --- | --- | --- | --- |
| cleaved Caspase 3 | unconjugated | Cell Signaling | 1679664S | 1/100 |
| CD31 | - | R&D | AF3628 | 1/200 |
| Donkey anti-Goat | Alexa Fluor 488 | Thermo Fisher | A11055 | 1/400 |
| Donkey anti-Goat | Alexa Fluor 555 | Thermo Fisher | A21432 | 1/400 |
| Donkey anti-Goat | Alexa Fluor 647 | Thermo Fisher | A21447 | 1/400 |
| Donkey anti-Rabbit | Alexa Fluor 488 | Thermo Fisher | A21206 | 1/400 |
| Donkey anti-Rabbit | Alexa Fluor 568 | Thermo Fisher | A10042 | 1/400 |
| Donkey anti-Rabbit | Alexa Fluor 647 | Thermo Fisher | A31573 | 1/400 |
| Donkey anti-Rat | Alexa Fluor 647 | Thermo Fisher | A21434 | 1/400 |
| Desmin | unconjugated | Abcam | ab15200 | 1/100 |
| ERG | unconjugated | Abcam | ab92513 | 1/200 |
| ERG | Alexa Fluor 647 | Abcam | ab196149 | 1/75 |
| GFP | FITC | Abcam | ab6662 | 1/400 |
| Golph4 | unconjugated | Abcam | ab28049 | 1/400 |
| pHistone H3 | Alexa Fluor 555 | Cell Signaling | 3475 | 1/100 |
| ICAM2 | unconjugated | BD Biosciences | 553326 | 1/100 |
| KLF4 | unconjugated | R&D | AF3158 | 1/20 |
| mCherry | Alexa Fluor 594 | Alfagene | M11240 | 1/100 |
| NG2 | unconjugated | Merck Millipore | AB5320 | 1/100 |
| pSmad1/5/9 | unconjugated | Cell Signaling | 13820 | 1/100 |
| αSMA | Cy3 | Sigma Aldrich | C6198 | 1/400 |

Tile-scan spanning of retinas were acquired on either a Zeiss Cell Observer Spinning Disk confocal microscope equipped with Zen blue software or a 3i Marianas SDC spinning disk confocal microscope equipped with SlideBook 6.0.22 software. An EC plan-neofluar Ph1 10x NA 0.30 dry objective, a plan-apochromat Ph2 20x NA 0.80 dry objective or an LD C-apochromat Corr 40x NA 1.10 water objective or a Plan-Apochromat DIC 40x NA 1.40 oil objective were used for the acquisitions. For EC volume, high-resolution images were obtained using a confocal laser point-scanning microscope (Zeiss 980) equipped with the Zen software, using a plan-fluor apochromat 63x NA 1.40 oil objective.

### Image analysis

Most of image analyses were performed with Fiji software^81^. Shunt occurrence was quantified manually as a ratio between the total AV shunt observed over the total AV sections quantified at a specific time point in a specific condition. Endothelial cell density was defined as the number of ERG+ or GFP+ (GNrep mouse strain) nuclei (quantified manually) over the CD31 corresponding surface (determined after thresholding based on fluorescence intensity). Endothelial proliferation was quantified manually as a ratio of the number of phospho-Histone H3/ERG or GFP double positive cells over the corresponding number of ERG or GFP+ cells. Diameters were quantified using the VasoMetrics Fiji tool^82^ with a 10 μm step between crosslines. Mural cell coverage was quantified as a surface ratio of each marker with CD31 surface after thresholding based on fluorescence intensity. Polarity was defined manually based on the angle of nucleus-to-Golgi axis with estimated flow direction in three categories: with the flow (0-45°), random (45°-135°), and against the flow (135°-180°). Sprouts were quantified manually as the ration of branch point number to a specific vessel (AV shunt or vein) over the length of that vessel. Neovascularised area was defined as the CD31 surface (after thresholding based on fluorescence intensity) over the proximal surface delimited by the AV shunt, its corresponding artery and vein, and the optic nerve. Lectin perfusion was quantified in two different ways. For neovascular area perfusion, it was defined as the lectin signal surface (after thresholding based on fluorescence intensity) over the proximal surface. For connection perfusion, it was quantified as a ration between the number of branch point positive for lectin and the total number of branch point considered.

Cell volume was analysed using Imaris software (Oxford Instruments). High-resolution images of single mGFP+ endothelial cells were acquired using a 980 confocal microscope equipped with a 63X NA 1.40 oil objective. The mGFP channel was segmented using “surfaces” segmentation tool to create a solid volume (sup.video 1). Unsegmented signal arising from the low GFP signal at the nucleus was manually corrected using Fiji.

### Statistical analysis

All statistical analyses were performed using R Studio (R version 1.4.1717, R Foundation for Statistical Computing). Quantifications were done on independent samples. Each data point corresponds to a shunt or a single cell, and the number of animals per experiment, as well as the number of litters, are stated in the figure legend. Statistical details of experiments are reported in the figures and their legends. No inclusion, exclusion or randomization criteria were used and all analysed samples were included. Comparisons between two experimental groups were analysed using Mann-Whitney test. Multiple comparisons between more than two experimental groups were assessed with Kruskal Wallis test and combined with Dunn post-hoc test using Benjamini & Hochberg correction for p-value adjustment. Proportion comparisons were analysed using Fisher exact t-test and combined with Fisher post-hoc test using Benjamini & Hochberg correction for p-value adjustment in the case of multiple comparisons. A result was considered significant when p < 0.05.

## Acknowledgements

We thank the VML lab and Fondation Leducq ATTRACT members for their helpful discussions. We thank IMM bioimaging and Rodent Facilities for outstanding support. This work was supported by European Research Council (679368), Fundação para a Ciência e Tecnologia (PTDC/MED-ANM/7695/2020 and EXPL/MED-ANM/1616/2021; CEECIND/04251/2017), Fondation LeDucq (17CVD03), European Commission (801423),

La Caixa Foundation (LCF/PR/HR22/52420027), and EU MSCA (842498).

## Author Contributions

Conceptualization: MO, AP, DR, SPO, CAF

Methodology: MO, AP, DR, NVC, TC, SPO, CAF

Experimentation and data acquisition: MO, AP, DR, NVC, TC, AF, MPS, YC, LHM, CAF

Data analysis: MO, AP, DR, NVC, TC, CAF

Project administration: CAF

Writing – original draft: MO, CAF

Writing – review & editing: MO, AP, DR, NVC, SPO, CAF

---

**Supplementary Figure 1.**
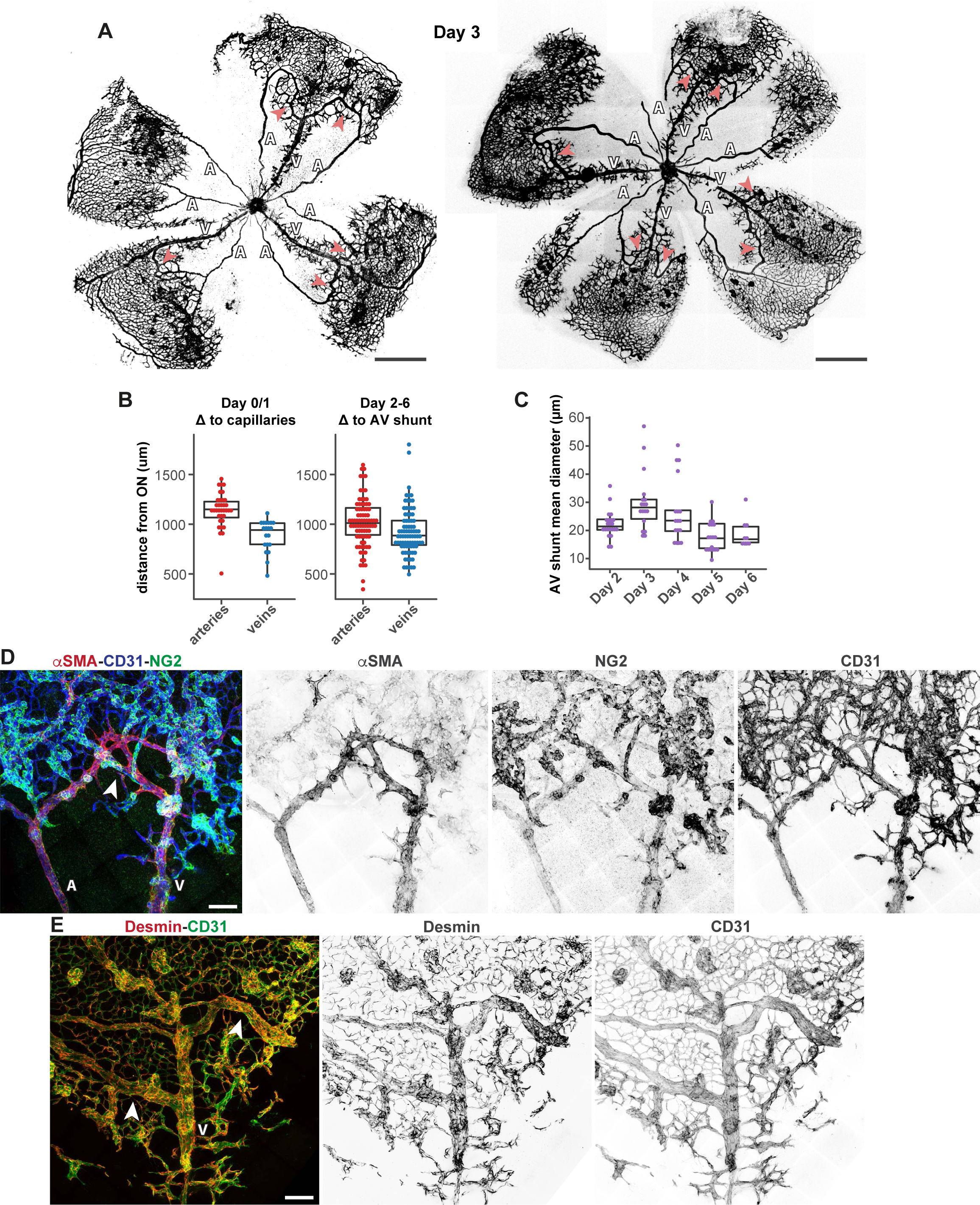
Characterization of OIR-induced AV shunts. **A,** Representative overview images of Day 3 mouse retinas stained for CD31 (grey). Red arrows: AV shunt; A: artery; V: vein. Scale bar: 500 µm. **B,** Quantification distance from optic nerve to capillaries (Day 0 and Day 1, left) or AV shunt (Days 2 to 6, right) on arteries or veins. Each dot represents an AV section from Day 0 and Day 1 (8 pups); and Days 2 to 6 (16 pups). P-values (ns) from Mann-Whitney test. **C,** Quantification of AV shunt mean diameterretinas exposed between Day 2 and Day 6. Each dot represents an AV shunt from Day 2 (4 pups); Day 3 (3 pups); Day 4 (3 pups); Day 5 (4 pups); and Day 6 (2 pups). P-values from Kruskal Wallis test and Dunn post-hoc test using Benjamini & Hochberg correction for multiple comparisons. **D,** Representative image of the vascular network stained for ECs (CD31, blue) and mural cells (αSMA, red; NG2, green) from a Day 4 mouse retina. Scale bar: 100 µm. **E,** Representative image of the vascular network stained for ECs (CD31, blue) mural cell coverage (desmin, red) from a Day 4 mouse retina. Scale bar: 100 µm.

**Supplementary Figure 2.**
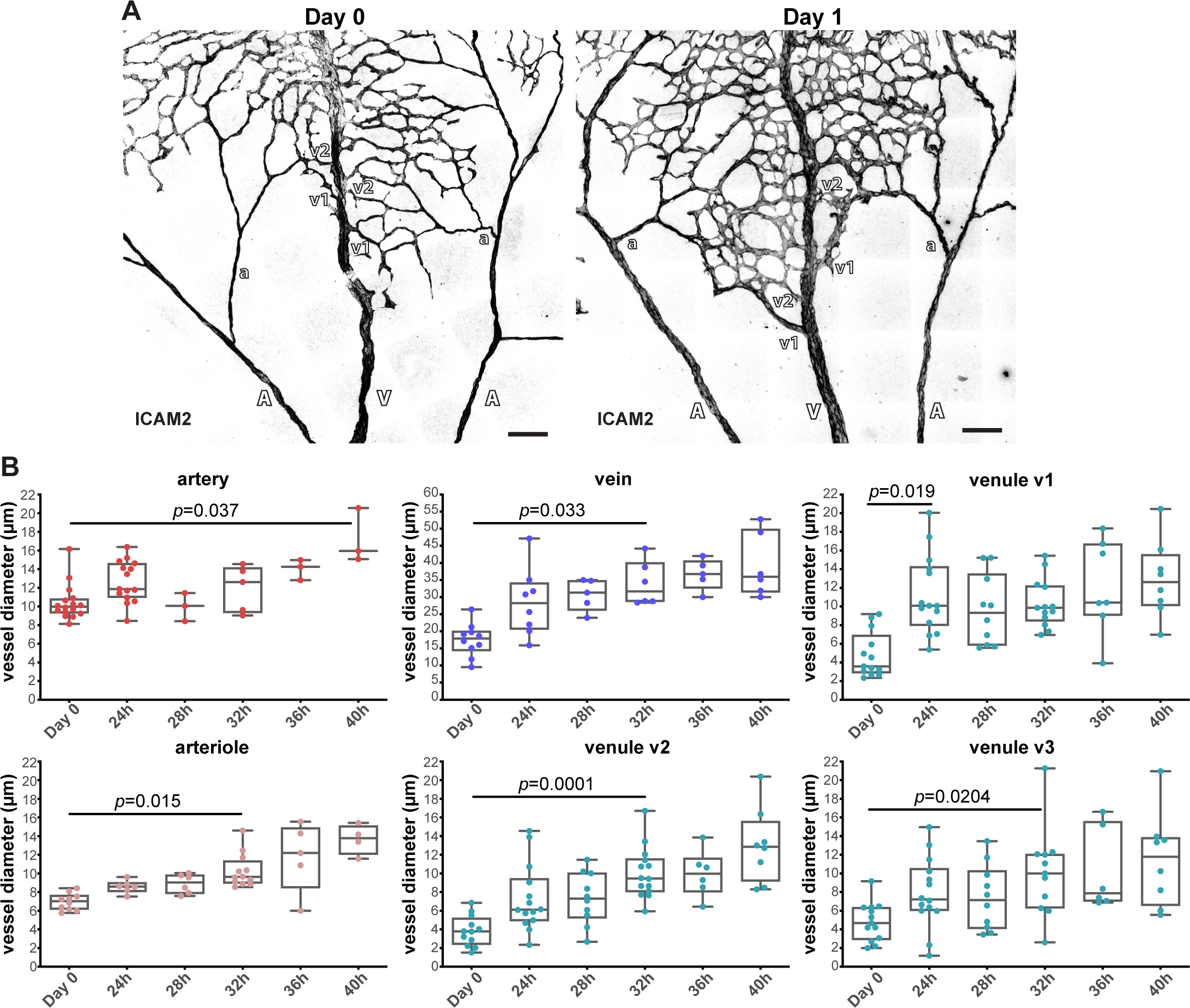
Asymmetric increased vessel diameter occurs prior to OIR-induced AV shunts. **A,** Representative images of mouse retinas stained for EC apical membrane (ICAM2, grey) at Day 0 and Day 1. A: artery; V: vein; a: arteriole; v: venule. Scale bar: 200 µm. **B,** Quantification of vessel diameter of arteries (left) and veins (right) from mouse retinas collected between Day 0 and 40h after returning to normoxia. Each dot represents a vessel from Day 0 (4 pups); 24h (4 pups); 28h (2 pups); 32h (4 pups); 36h (3 pups); and 40h (2 pups). P-value from Krustal-Wallis test with Dunn’s correction for multiple comparisons.

**Supplementary Figure 3.**
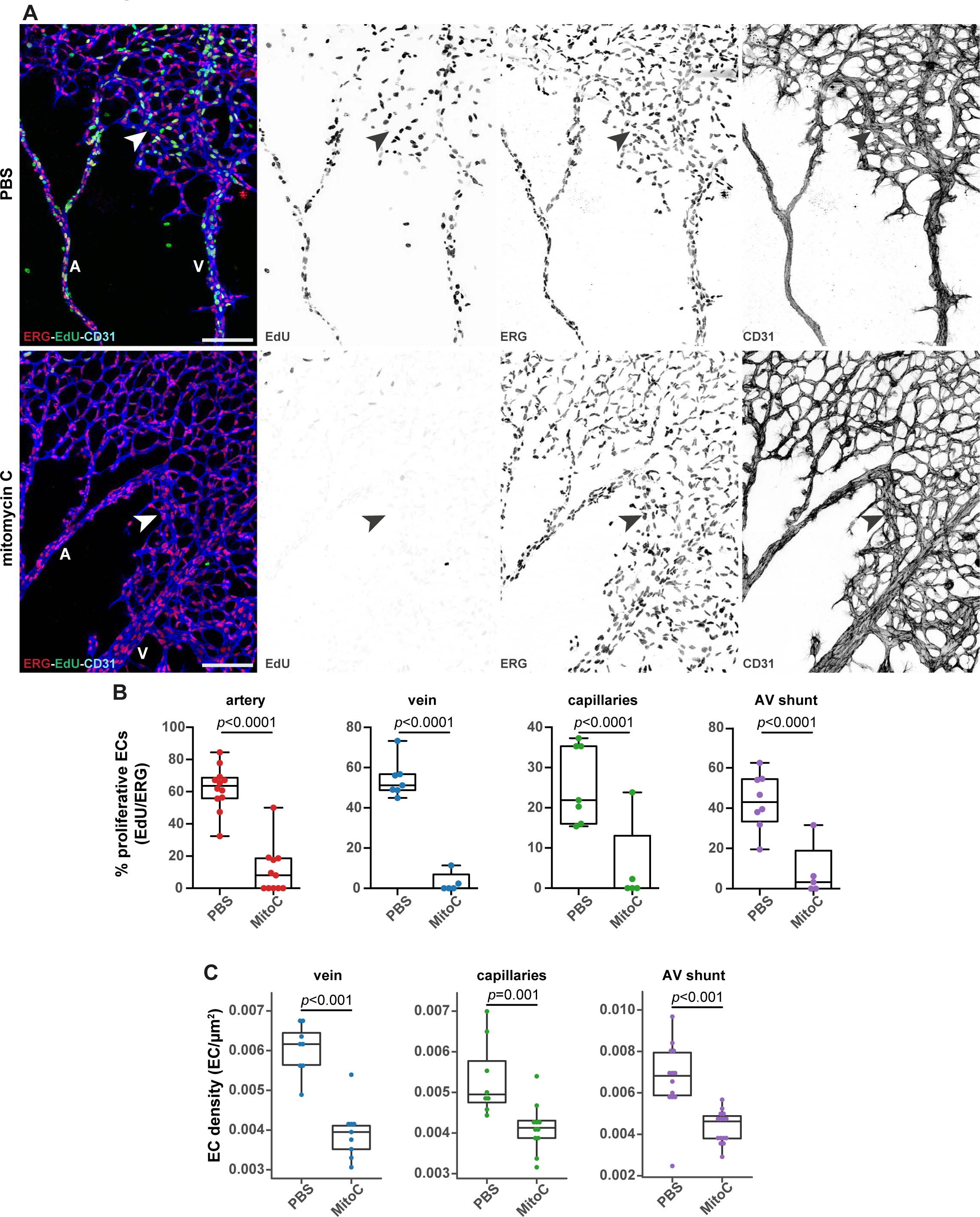
OIR-induced AV shunts form independently of EC proliferation. **A,** Representative images of Day 2 mouse retinas treated with PBS or MitoC stained for EC nuclei (ERG, red), proliferative cells (EdU, green) and EC membrane (CD31, blue). Arrowhead: AV shunt; A: artery; V: vein. Scale bar: 200 µm. **B,** Quantification of proliferative ECs (EdU/ERG ratio) in AV shunts at Day 2. Each dot represents a specific vessel bed from different retinas. P-values from Mann-Whitney test. **C,** Quantification of EC density in veins, capillaries and AV shunts from Day 3 retinas treated with PBS (4 pups) and mitomycin C (5 pups). Each dot represents one vessel. P-values from Mann-Whitney test.

**Supplementary Figure 4.**
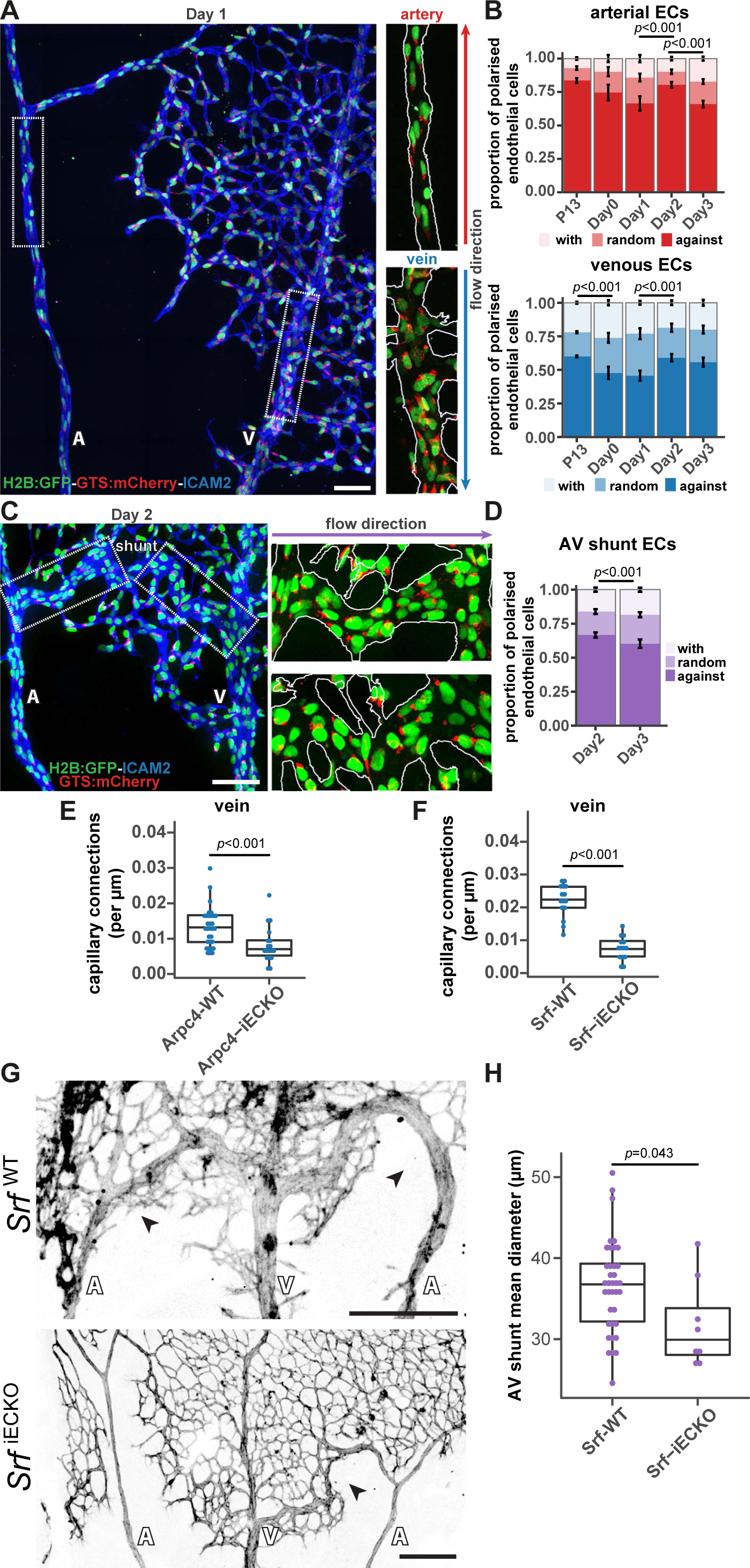
Neither EC polarity nor EC migration is significantly associated with AV shunt development. **A,** Representative image of a Day 1 GNrep mouse retina stained for ICAM2 (blue). EC nuclei are labelled by nGFP (green) and EC Golgi apparatuses are labelled by mCherry (red). A: artery; V: vein. Scale bar: 100 µm. Magnifications of an artery (top) and a vein (bottom). Red/blue arrows: blood flow direction in artery and vein, respectively; dotted white lines: vessel outline based on ICAM2 staining. **B,** Quantification of EC polarization pattern in arteries (top) and veins (bottom) between Day 0 and Day 3 retinas (3 pups for all conditions) and in non-OIR retinas (5 pups). P-values from Fisher exact t-test and Fisher post-hoc test using Benjamini & Hochberg correction for multiple comparisons. **C,** Representative image of a Day 2 GNrep mouse retina stained for ICAM2 (blue). EC nuclei are labelled by nGFP (green) and EC Golgi apparatuses are labelled by mCherry (red). A: artery; V: vein. Scale bar: 100 µm. Highlights of AV shunt regions. Purple arrows blood flow direction; dotted white lines: vessel outline based on ICAM2 staining. **D,** Quantification of EC polarization proportions in AV shunt at Day 2 and Day 3 (3 pups). P-value from Fisher exact t-test and Fisher post-hoc test using Benjamini & Hochberg correction for multiple comparisons. **E,** Quantification of vein to neovascular capillary connections per µm in Day 3 Arpc4-WT (7 pups) and Arpc4-iECKO (5 pups) mouse retinas. Each dot represents a vein. P-value from Mann-Whitney test. **F,** Quantification of vein to neovascular capillary connections per µm in Day 3 Srf-WT (8 pups) and Srf-iECKO (5 pups) mouse retinas. Each dot represents a vein. P-value from Mann-Whitney test. **G,** Representative images of Day 3 Srf-WT (top) and Srf-iECKO (bottom) retinas stained for ECs (CD31, grey). Black arrows: AV shunts; A: artery; V: vein. Scale bar: 100 µm. **H,** Quantification of AV shunt mean diameter in Day 3 in Srf-WT (6 pups) and Srf-iECKO (3 pups) retinas. Each dot represents an AV shunt. P-value from Mann-Whitney test.

**Supplementary Figure 5.**
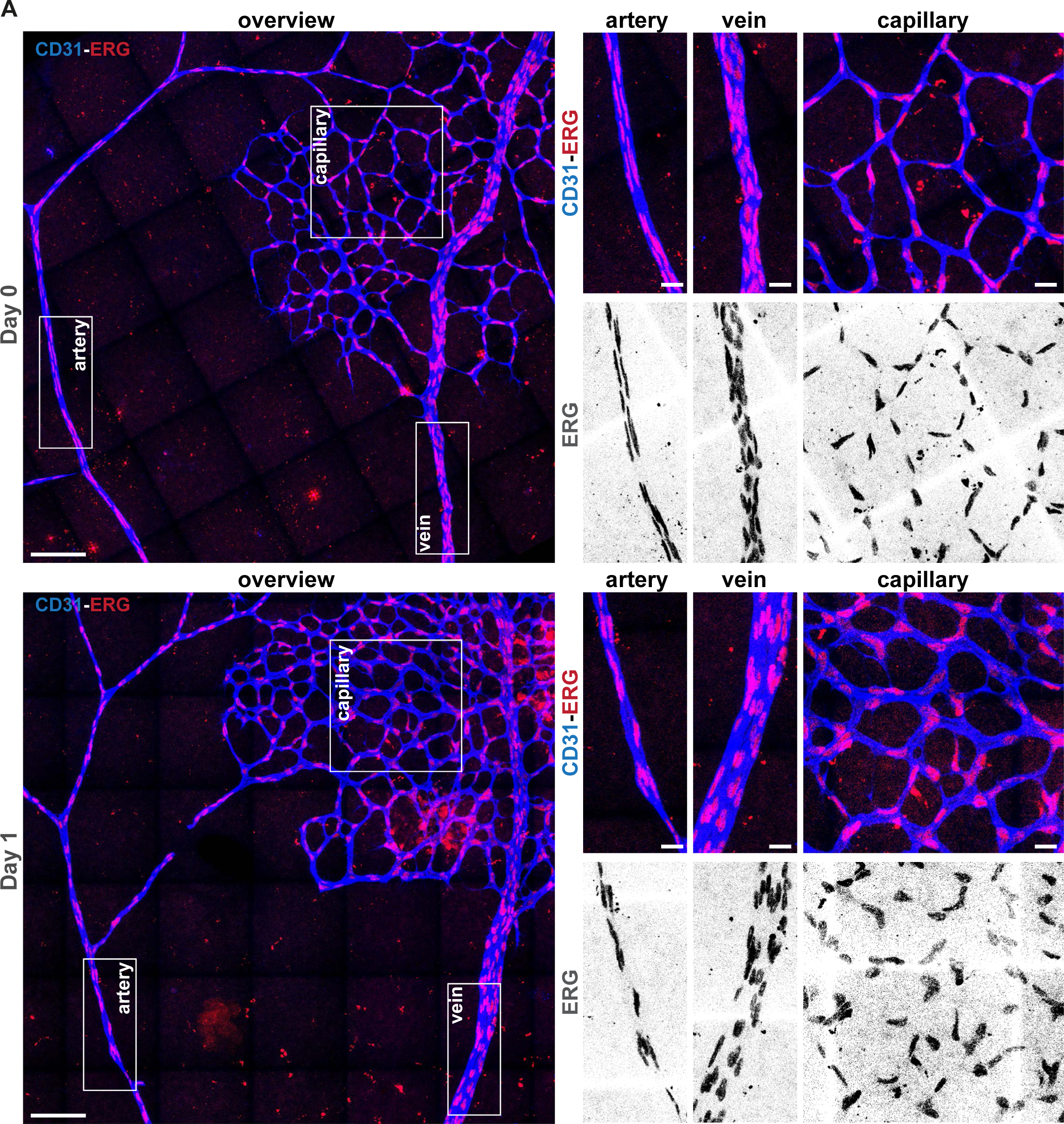
A decrease in EC density precedes AV shunt formation. **A,** Representative images of Day 0 (top panel) and Day 1 (bottom panel) retinas stained for EC nuclei (ERG, red/grey) and EC membrane (CD31, blue) highlighting a decrease in EC density. Scale bar: 100 µm. Magnifications of arterial, venous and capillary regions.

**Supplementary Figure 6.**
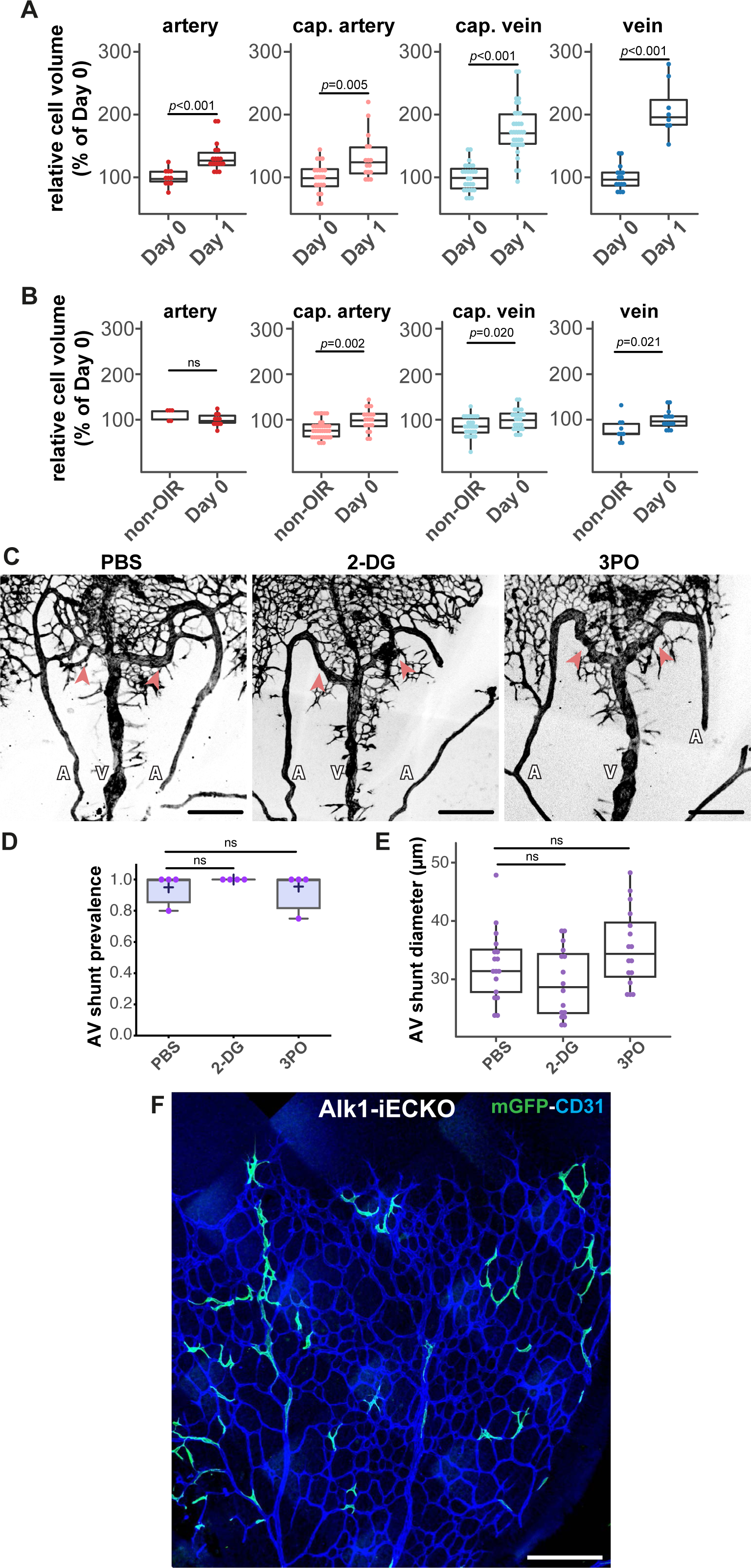
Inhibition of glycolysis does not alter EC volumes nor AV shunt formation. **A,** Quantification of normalized EC volume (% mean EC volume at Day 0) in arteries, arterial capillaries, venous capillaries, and veins from Day 0 and Day 1. Each dot represents one EC from Day 0 (3 pups) and Day 1 (3 pups). P-values from Mann-Whitney test. **B,** Quantification of normalized EC volume (% mean cell volume in non-OIR) in arteries, arterial capillaries, venous capillaries, and veins from non-OIR and Day 0 retinas. Each dot represents one EC from non-OIR (2 pups) and Day 0 (3 pups). P-values from Mann-Whitney test. **C,** Representative images of Day 3 retinas treated with PBS or 2-DG or 3PO stained for ECs (CD31, grey). Red arrows: AV shunts; A: artery; V: vein. Scale bar: 100 µm. **D,** Quantification of AV shunt prevalence at Day 3 retinas treated with PBS (24 AV sections, 2 pups), 2-DG (18 AV sections, 2 pups), and 3PO (22 AV sections, 2 pups). P-values (ns) from Kruskal Wallis test and Dunn post-hoc test. **E,** Quantification of AV shunt mean diameter at Day 3 in retinas treated with PBS (2 pups), 2-DG (2 pups), and 3-PO (2 pups). Each dot represents an AV shunt. P-values from Kruskal Wallis test and Dunńs correction for multiple comparisons. **F,** Representative image of a P6 Alk1-iECKO-mTmG retina iECKO-mTmG 24h post-tamoxifen injection highlighting mosaic activation of mGFP (green) and co-stained for EC membrane (CD31, blue) Scale bar: 200 µm.

**Supplementary Figure 7.**
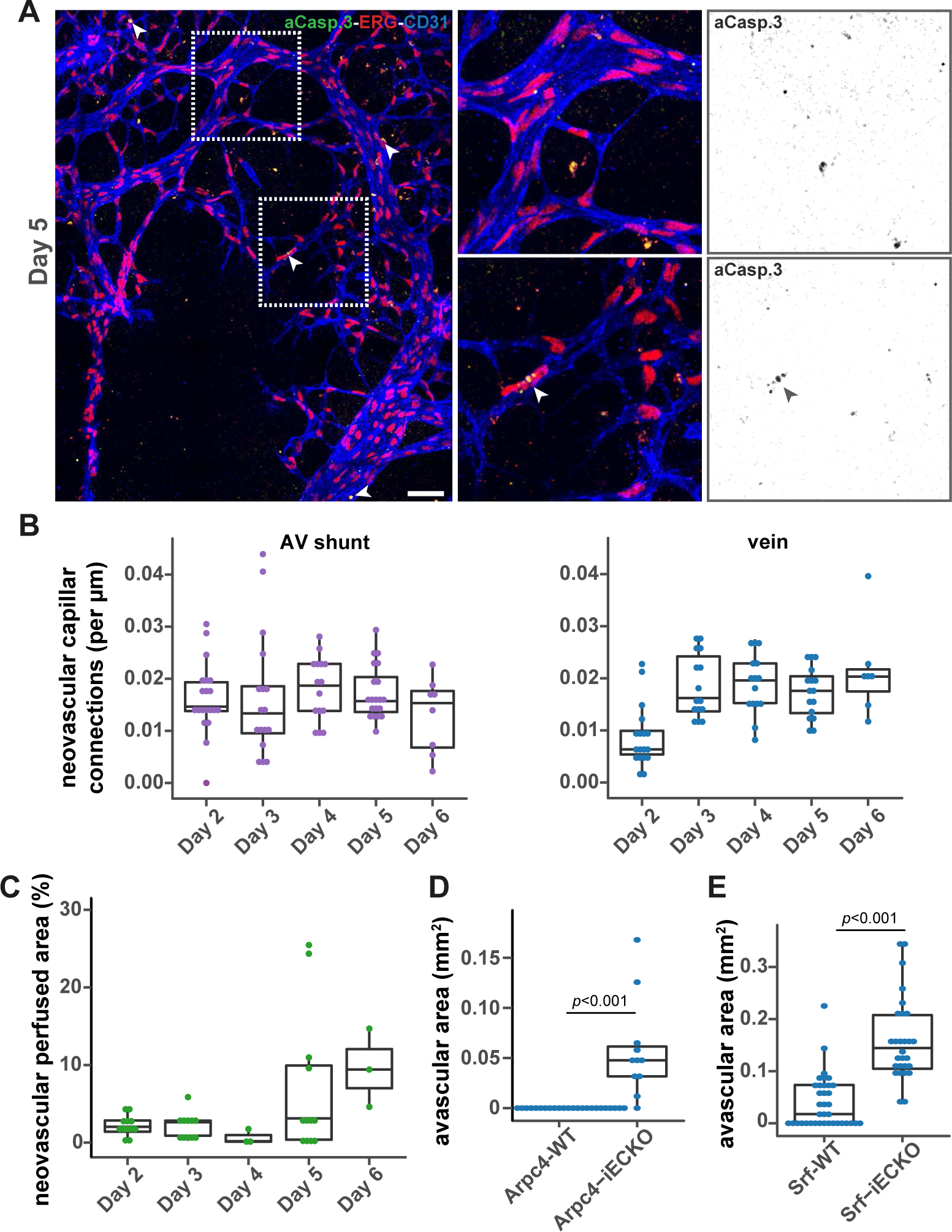
AV shunt regression does not involve EC apoptosis but requires neo-angiogenesis. **A,** Representative image of a Day 5 retina stained for EC membrane (CD31, blue), EC nuclei (ERG, red) and apoptotic cells (active cleaved caspase 3, green). White arrows: AV shunts; A: artery; V: vein; red arrow: apoptotic EC. Scale bar: 100 µm. **B,** Quantification neovascular capillary connections arising from AV shunts (left) or veins (right) per µm from mouse retinas collected between Day 2 and Day 6. Each dot represents an AV shunt or a vein from Day 2 (4 pups); Day 3 (3 pups); Day 4 (3 pups); Day 5 (5 pups); and Day 6 (2 pups). P-values from Kruskal Wallis test and Dunn post-hoc test using Benjamini & Hochberg correction for multiple comparisons. **C,** Quantification of perfused neovascular capillary area (% of the proximal area) between Day 2 and Day 6. Each dot represents a proximal region from Day 2 (6 retinas); Day 3 (4 retinas); Day 4 (1 retina); Day 5 (5 retinas); and Day 6 (3 retinas). **D,** Quantification of the avascular area in Day 7 mouse retinas from in Arpc4-WT (4 pups) and Arpc4-iECKO (4 pups). Each dot represents a proximal AV shunt region. P-value from Mann-Whitney test. **E,** Quantification of the avascular area in Day 9 mouse retinas from Srf-WT (9 pups) and Srf-iECKO (7 pups). Each dot represents a proximal AV shunt region. P-value from Mann-Whitney test.

**Supplementary Video 1 – Example of EC volume segmentation.** 3D rotational movie showing an example of a segmented single EC (yellow) from of a D1 Cdh5-CreERT::R26-mTmG retina. Retinas were stained for EC membrane (CD31, red) and EC nuclei (ERG, blue), and endogenous membrabar GFP signal is shown (green).

